# AI-enabled discovery of small molecules targeting complementary pathways for hair follicle rejuvenation

**DOI:** 10.64898/2026.06.09.728282

**Authors:** Zihan Qu, Yang Li, Stella E Cho, Lara Doğan, Qing Yao, Lan Tang, Gary Zhao, Evan M Zhao, Alicia Li, Satotaka Omori, Felix Wong, David KY Zhang

## Abstract

Hair thinning arises from multi-faceted dysfunction within the hair follicle, driven by both intrinsic cellular pathways and pathways responding to extrinsic hormonal and microenvironmental cues. Here, we present an AI-enabled discovery framework to discover small molecules that promote hair follicle rejuvenation. This framework integrates graph neural networks trained on phenotypic screening data with structure-based virtual screening to prioritize compounds that modulate complementary biological pathways. Through AI-enabled screening, hit-to-lead optimization, and medicinal chemistry, we identified four compounds that increase follicle dermal papilla cell viability, stabilize hypoxia signaling by inhibiting prolyl hydroxylase domain protein 2 (PHD2), and suppress androgen-mediated follicular miniaturization by inhibiting 5α-reductases (5-ARs). RNA sequencing analyses confirmed pathway engagement, and functional validation across primary cells and a 3D hair follicle organoid model demonstrated high activity and cellular specificity. The lead compounds were incorporated into a water-based formulation, where they demonstrated robust solubility and combinatorial efficacy to increase sprouting length of follicle organoids. These results establish an AI-enabled platform for discovering multi-pathway modulators of hair follicle rejuvenation.

## Introduction

Hair thinning is a highly prevalent condition affecting both men and women. Beyond its visible effects, hair shedding, thinning, and ultimately, hair loss can cause substantial psychological burden among individuals^1^. Despite its impact, modern solutions remain limited. Approved treatments are repurposed drugs that have suboptimal efficacy and significant adverse effects. Minoxidil, initially discovered as a vasodilator^2^, promotes hair growth but elicits inconsistent and suboptimal responses^3,4^. Complications such as rapid hair shedding during early usage have been reported^5^. On the other hand, finasteride and its more potent derivative dutasteride, developed as benign prostatic hyperplasia drugs^6,7^, promote hair growth in men but may induce reduced libido, erectile dysfunction and gynecomastia^8^. Alternative solutions, such as platelet-rich plasma, low-level red laser therapy, microneedling, and hair transplantation, are also frequently used in clinical practice^3^, but their efficacy is often variable, and they can elicit side effects such as scarring^9,10^.

Progress in developing new treatments has been slow. Several candidates in clinical testing continue to rely on repurposed drugs, including clascoterone (a steroidal androgen-receptor antagonist first developed for acne)^11^, TH07 (a regimen incorporating minoxidil, finasteride, and latanoprost, a prostaglandin F2α analog originally developed for glaucoma)^12^, and topical cetirizine (an antihistamine thought to modulate prostaglandin signaling in the follicular microenvironment)^13^. Only a few emerging therapies target new mechanisms, such as PP405, a topical mitochondrial pyruvate carrier inhibitor designed to reactivate dormant hair-follicle stem cells with reported stimulation of vascular endothelial growth factor (VEGF) and lactate production^14^, and ABS-201, an investigational anti-prolactin receptor biologic intended to preserve the stem/progenitor niche^15^. In parallel, the lack of safe and effective therapeutics has expanded a large non-drug market based on supplements such as flavonoids, but many supplements have limited target specificity, weak mechanistic support or poor chemical stability.

Another major bottleneck is that most interventions address isolated pathways within a highly complex hair follicle system. Hair follicle rejuvenation depends on multiple intrinsic and extrinsic programs^16^. At the intrinsic level, follicle dermal papilla cells (DPCs) regulate hair follicle size, cycling, and regenerative capacity. With age, stress, and environmental perturbation, DPCs lose growth potential, metabolic activity, and inductive function^17^. At the extrinsic level, the surrounding niche provides key metabolic, vascular, and hormonal signals to support the hair follicle. Hypoxia-inducible factor-1α (HIF-1α) is one such upstream regulator of major regenerative programs in the follicular microenvironment, driving angiogenic signaling, metabolic adaptation, and tissue-protective responses^18^. Prolyl hydroxylase domain protein 2 (PHD2) inhibition stabilizes HIF-1α and activates downstream regenerative programs, such as VEGF and lactate production^19^. Canonical PHD2 inhibitors such as L-mimosine have been shown to stabilize HIF-1α and promote downstream angiogenesis and regenerative responses^20^. In parallel, androgen signaling remains a major extrinsic driver of hair follicle miniaturization. 5α-reductases (5-ARs) catalyze the conversion of testosterone to DHT, which accumulates within the scalp^21^. This activates androgen receptor signaling, thereby shortening the anagen phase, prolonging the telogen phase, progressively driving follicle miniaturization^22^.

Artificial intelligence (AI) is increasingly used in small-molecule discovery to search, rank, and prioritize chemical spaces by narrowing large libraries into smaller and experimentally tractable compound sets. Building on this principle, we developed an AI-enabled discovery framework to address hair follicle rejuvenation by integrating the cell-intrinsic phenotype of DPC viability with two cell-extrinsic pathways involving PHD2 and 5-ARs. Our framework led to the discovery of four compounds: NV-623 and NV-624, which enhanced DPC proliferation and metabolic activity; NV-273, which inhibited PHD2 (inducing durable HIF-1α pathway activation); and NV-1065, a non-steroidal 5-AR inhibitor with comparable potency as dutasteride and no cellular androgen receptor activity. In hair follicle organoids, NV-623 and NV-624 treatments increased organoid sprouting length, while NV-1065 rescued testosterone-induced growth inhibition. In primary human cells, NV-273 induced VEGF and lactate production. Medicinal chemistry improved the solubility of the lead compounds and enabled their incorporation into a water-based formulation. In the formulation, the compounds retained their activity and showed combinatorial efficacy in promoting follicle organoid elongation. Together, these results establish an AI-enabled and mechanistically grounded strategy for new molecule development.

## Results

### AI-enabled discovery of multi-pathway modulators for hair follicle rejuvenation

We developed an AI-guided discovery framework integrating the intrinsic DPC phenotype with extrinsic regulatory pathways involving PHD2 and 5-ARs (Figs. 1a, b). For the phenotype-aware DPC viability screening, we used an ensemble-based graph neural network pipeline trained on empirical screening data, as previously described^23–25^. An initial library of 50,000 chemically diverse compounds was curated based on structural diversity, molecular weight, and exclusion of known toxic or highly reactive chemotypes. Primary human DPCs derived from two independent donors were cultured under supplier-recommended conditions, and changes in cell viability were measured by CellTiter-Glo^®^ (CTG) and CellTiter-Blue^®^ (CTB). Robotic high-throughput screening (HTS) generated a training dataset for supervised learning, in which an ensemble of ten Chemprop models was trained to predict phenotype–molecular structure relationships^23^. The trained ensemble was subsequently deployed to screen 11,278,376 compounds compiled from publicly available and commercially accessible small-molecule databases, yielding 22,506 compounds (0.20%) with prediction scores greater than 0.032 (Fig. 1c). Application of additional constraints, including molecular weight range of 220–475 Da and structural liability filters (PAINS, Brenk, Lipinski, and steroidal filters), reduced the candidate set to 2,029 compounds. Finally, 157 compounds were prioritized for screening, and 33 hits were identified in the initial phenotypic assay, representing a 21.0% hit rate.

**Figure 1.**
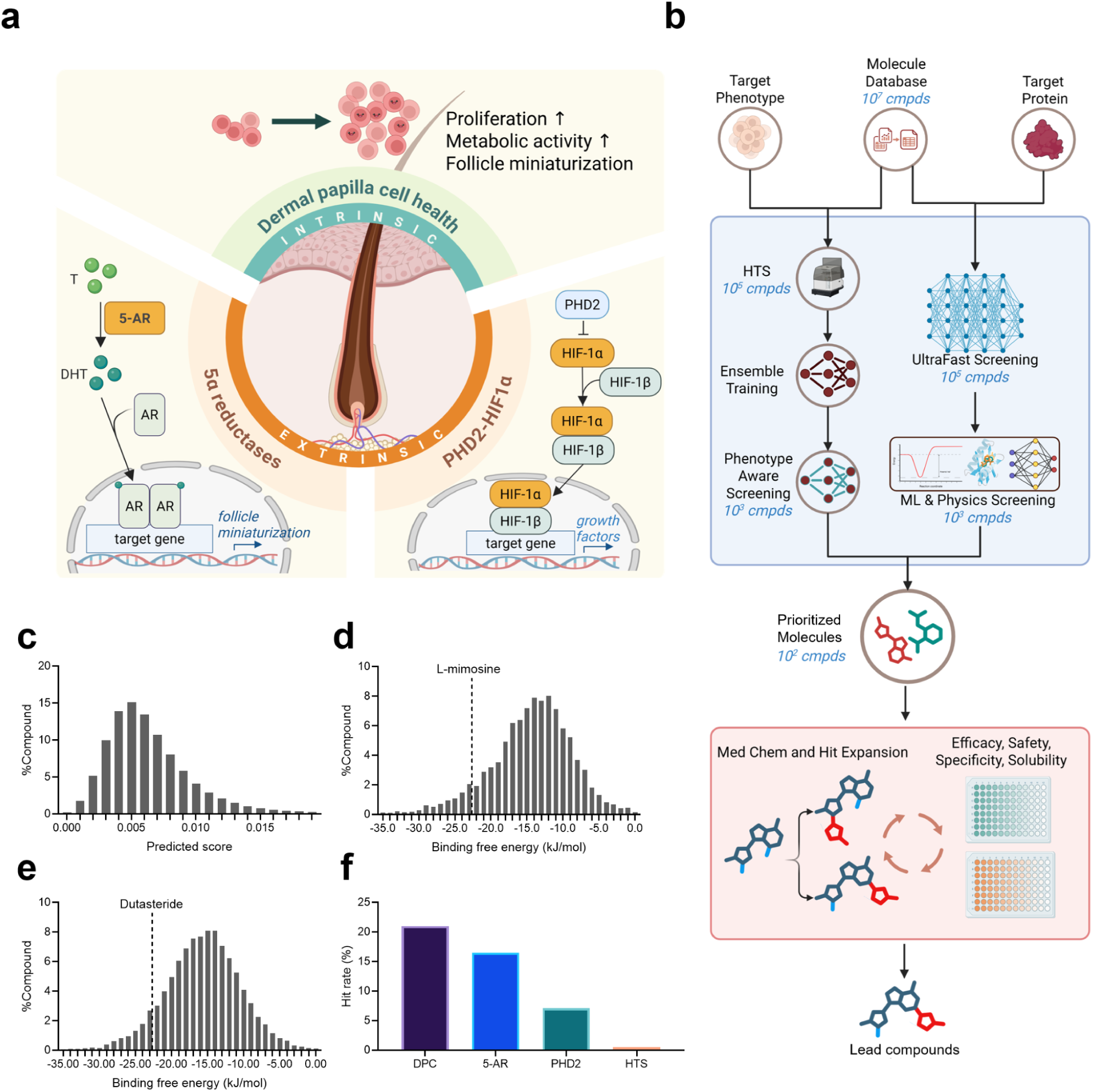
AI-enabled discovery of small-molecule modulators targeting complementary pathways in hair thinning and rejuvenation. (a) Schematic representation of complementary pathways regulating hair follicle rejuvenation. Intrinsic pathways are governed by DPC viability, while extrinsic pathways are governed by the PHD2-HIF-1α axis and 5α-reductases. (b) AI-enabled virtual screening and experimental validation framework. Libraries of over 10 million compounds were processed through sequential computational filtering, molecular representation learning, and structure–property optimization to discover candidate compounds for *in vitro* validation. Rank-ordered score distributions from virtual screening for (c) DPC proliferation modulators, (d) PHD2 inhibitors, and (e) 5-AR inhibitors. (f) Substantial improvement in hit rate of screening compounds prioritized by our computational framework.

In parallel, we implemented a multi-stage computational screening and prioritization framework targeting extrinsic regulators of the hair follicle. Target-aware predictive models were combined with physics-based approaches, including Poisson–Boltzmann surface area (PBSA) calculations, to prioritize candidates based on predicted binding affinity. Structure–property optimization was then applied to refine compounds with respect to potency, safety, and solubility (Fig. 1b). Rank-ordered score distributions from the virtual screenings revealed strong enrichment of high-confidence candidates for both PHD2 and 5-AR inhibition (Figs. 1c-e). Using L-mimosine and dutasteride as benchmark compounds for PHD2 and 5-AR inhibition, respectively, we selected the top 2.0% of ranked compounds, corresponding to 1,953 PHD2 inhibitor candidates and 710 5-AR inhibitor candidates. For PHD2 inhibition, 210 compounds were prioritized for screening, and 7.1% of these compounds exhibited inhibitory activity against PHD2. For 5-AR inhibition, 406 compounds were prioritized for screening, of which 16.5% of the prioritized compounds exhibited inhibitory activity against 5-AR. Our AI-guided discovery framework improved hit rates relative to conventional HTS, achieving hit rates ranging from 7.1% to 21.0%, depending on target class. This represents an over 100-fold hit rate increase compared with the ∼0.10% to 1% hit rates typically observed in traditional HTS campaigns^26–28^.

### Medicinal chemistry yielded DPC potentiators NV-623 and NV-624

Primary DPC donors were selected to represent an age range most commonly affected by hair thinning (Supplementary Table 1). DPCs were treated with the computationally prioritized compounds, and changes in cell proliferation were measured using CTG (Fig. 2a). CTG responses were normalized to vehicle-treated cells and compounds exhibiting greater than 5.0% enhancement of DPC proliferation were prioritized for dose–response validation. A nonspecific corticoid, prednisolone, was used as the assay benchmark for DPC proliferation^29,30^.

**Figure 2.**
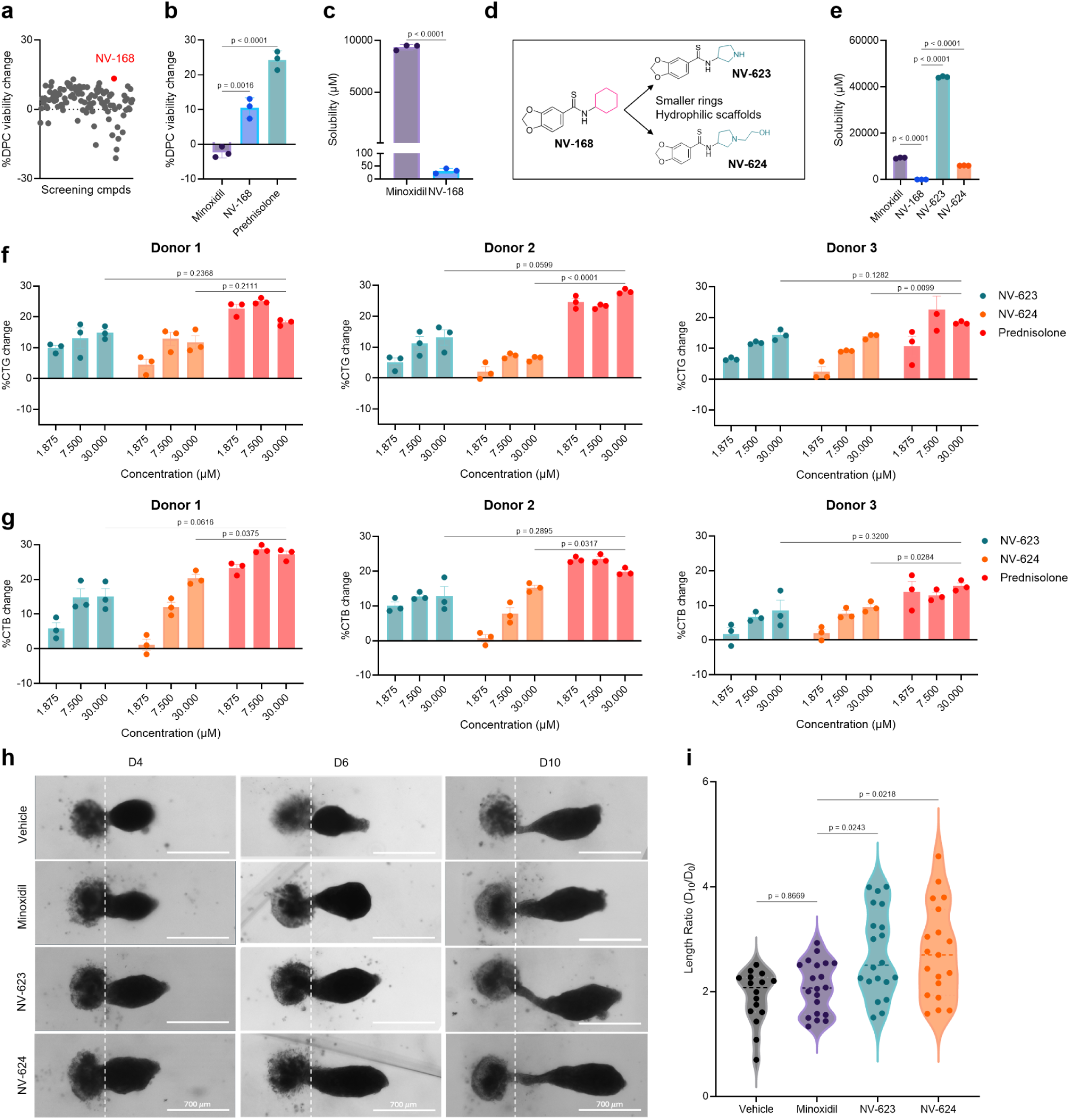
NV-623 and NV-624 promoted DPC proliferation and hair follicle organoid sprouting. (a) Summary of normalized changes in CTG responses in DPCs for compounds prioritized through virtual screening. (b) NV-168 promoted greater CTG responses than minoxidil. CTG changes were determined by normalizing to vehicle controls. (c) NV-168 was less soluble than minoxidil. (d) Medicinal chemistry strategy to improve NV-168 aqueous solubility yielded NV-623 and NV-624. (e) Both NV-623 and NV-624 showed higher aqueous solubility than the parent compound NV-168, and NV-623 showed higher aqueous solubility than the benchmark compound minoxidil. NV-623 and NV-624 retained efficacy in promoting DPC proliferation, as measured by (f) CTG and (g) CTB assays across three independent donors. (h) Representative images showing NV-623 and NV-624 promoted hair follicle organoid sprouting. The white dashed line marks the position from which the follicle organoid sprouting initiates. (i) Quantification of hair follicle organoid sprouting length ratio at day 10 relative to day 0. Data in (a) are representative of the mean of n=3 technical replicates from 3 experimental replicates. Data in (b-c), (e-g), and (i) represent mean ± s.d. from n=3 technical replicates and are representative of over 3 experimental replicates. Data in (h) are representative of n=16 (vehicle), n=20 (minoxidil and NV-623), and n=19 (NV-624) technical replicates from over 3 experimental replicates. p values for pairwise comparisons between indicated groups are shown above bars, calculated using one-way ANOVA followed by Tukey’s multiple comparisons test.

Among the confirmed hits, NV-168 emerged as a top candidate, increasing DPC proliferation by 11% relative to vehicle controls (Fig. 2b).

Despite its biological activity, NV-168 exhibited poor aqueous solubility. LC–MS-based measurements revealed a water solubility of roughly 32 μM, compared with 9,338 μM for minoxidil, a commonly used hair-growth agent (Fig. 2c). To improve its aqueous solubility, we conducted medicinal chemistry focusing on modifying NV-168’s terminal ring scaffold and substituents (Fig. 2d). This optimization yielded two lead compounds, NV-623 and NV-624, with substantially higher aqueous solubility. NV-624 exhibited an aqueous solubility of 7,196 μM, only moderately lower than that of minoxidil (p < 0.0001), while NV-623 achieved a higher solubility of 38,048 μM, representing a more than 2,000-fold improvement relative to the parent compound NV-168 (p < 0.0001, Fig. 2e).

Although certain structural modifications in NV-168 led to reduced efficacy, NV-623 and NV-624 consistently enhanced DPC viability across three independent donors in a dose-dependent manner as quantified by CTG and CTB (Figs. 2f, g; Supplementary Fig. 1a). The maximal proliferative response consistently exceeded 15% when normalized to vehicle control (0.1% DMSO) in all donors. This increase in DPC proliferation was further confirmed with cell nuclear staining and enumeration - NV-623 and NV-624 increased the cell count by over 15% relative to the vehicle control (Supplementary Figs. 1c, d).

Next, we evaluated the cellular selectivity and potential toxicity of NV-623 and NV-624 across a panel of primary skin and non-skin cells and human cell lines. At 30 μM, neither NV-623 nor NV-624 induced detectable primary macrophage activation or genotoxic transformation potential. By contrast, the positive controls elicited the expected responses, with 1 μg/mL of lipopolysaccharide (LPS) inducing ∼20% macrophage activation and 4-nitroquinoline 1-oxide (4NQO), a known tumorigenic compound, causing genotoxicity at 6.5 μM. (Supplementary Fig. 2a, b). Both NV-623 (2.6%) and NV-624 (1.6%) showed negligible ARE-Nrf2 pathway activation in HepG2 reporter cells, a common readout of electrophilic or oxidative stress responses, in contrast to quercetin (207%) and minoxidil (18%) especially at 30 μM (Supplementary Fig. 2c). Consistent with this result, both compounds produced negligible reactive oxygen species (ROS) production in primary DPCs and human epidermal keratinocytes (HEKs) across a 1.5-log concentration range, whereas quercetin induced around 20% ROS at all doses tested (Supplementary Fig. 2d). Across a panel of primary skin cells, both NV-623 and NV-624 were well tolerated as quantified by CTG signal changes. Both compounds selectively enhanced DPC viability with only mild toxicity observed from NV-624 at high concentrations in human dermal microvascular endothelial cells (HDMEC; Supplementary Fig. 2e). Across non-skin cell types, the compounds showed no significant changes to cell viability except for mild toxicity in human renal cortical epithelial cells at high concentrations (HRCEpC; NV-623 only, Supplementary Fig. 2f). In contrast to our NV-623 and NV-624, minoxidil and prednisolone exhibited reduced selectivity or greater toxicity (Supplementary Fig. 2e and f). Minoxidil showed clear toxicity in DPCs and also reduced viability in human dermal fibroblasts (HDF) and human primary preadipocytes (HPAd). Prednisolone, as expected, increased proliferation in human epidermal keratinocytes (HEKs), HDF, and HPAd in addition to DPCs. It also reduced the metabolic activity of peripheral blood mononuclear cells (PBMC) at higher doses as quantified by a reduction in the CTB response, consistent with its immunosuppressive functions (Supplementary Fig. 2f)^32^.

To evaluate NV-623 and NV-624 in a more physiologically relevant context, we leveraged a human hair follicle organoid model generated by co-culture of primary DPCs and HEKs (Supplementary Fig. 1d). This system recapitulates key epithelial–mesenchymal interactions required for follicular morphogenesis and growth^33^. Follicle organoids treated with NV-623 or NV-624 showed significantly greater sprouting length over a 10-day culture period compared to those treated with minoxidil or the vehicle (Fig. 2h). At day 10, NV-623 and NV-624 exhibited stronger efficacy than minoxidil in increasing follicle organoid sprouting length (Fig. 2i, p = 0.0243 for NV-623; p = 0.0218 for NV-624), consistent with their enhanced proliferative effects in primary DPC CTG/CTB assays. Together, these findings highlight NV-623 and NV-624 as efficacious and selective potentiators of human DPCs.

### NV-273 induced durable HIF-1α activation and downstream regenerative programs

We developed a cell-based hypoxia response element luciferase (HRE-Luc) reporter assay in HEK293T cells to screen for PHD2 inhibitors. In this system, PHD2 inhibition stabilizes HIF-1α, resulting in increased transcription of a luciferase reporter, which enables quantification of HIF-1α signaling (Fig. 3a). A set of reported PHD2 inhibitors was tested to validate the assay^34–36^, and L-mimosine demonstrated the most robust dose-dependent HIF-1α activation (Supplementary Fig. 3a). Although deferoxamine (DFOA) exhibited stronger effects, the lack of dose-dependency in this assay suggested potential nonspecific interactions^37,38^, rendering it a less appropriate benchmark than L-mimosine. Virtual screening–prioritized compounds were evaluated at 15 µM in this reporter system, and reporter activation was normalized to vehicle-treated controls (Fig. 3b). Notably, more than 30% of the tested compounds produced reporter activation greater than 10%, supporting the effectiveness of the computational prioritization. As a starting point to identify hits for further optimization, we used 30% HIF-1α activation as the threshold for hit selection to minimize noise and poor responders^39^.

**Figure 3.**
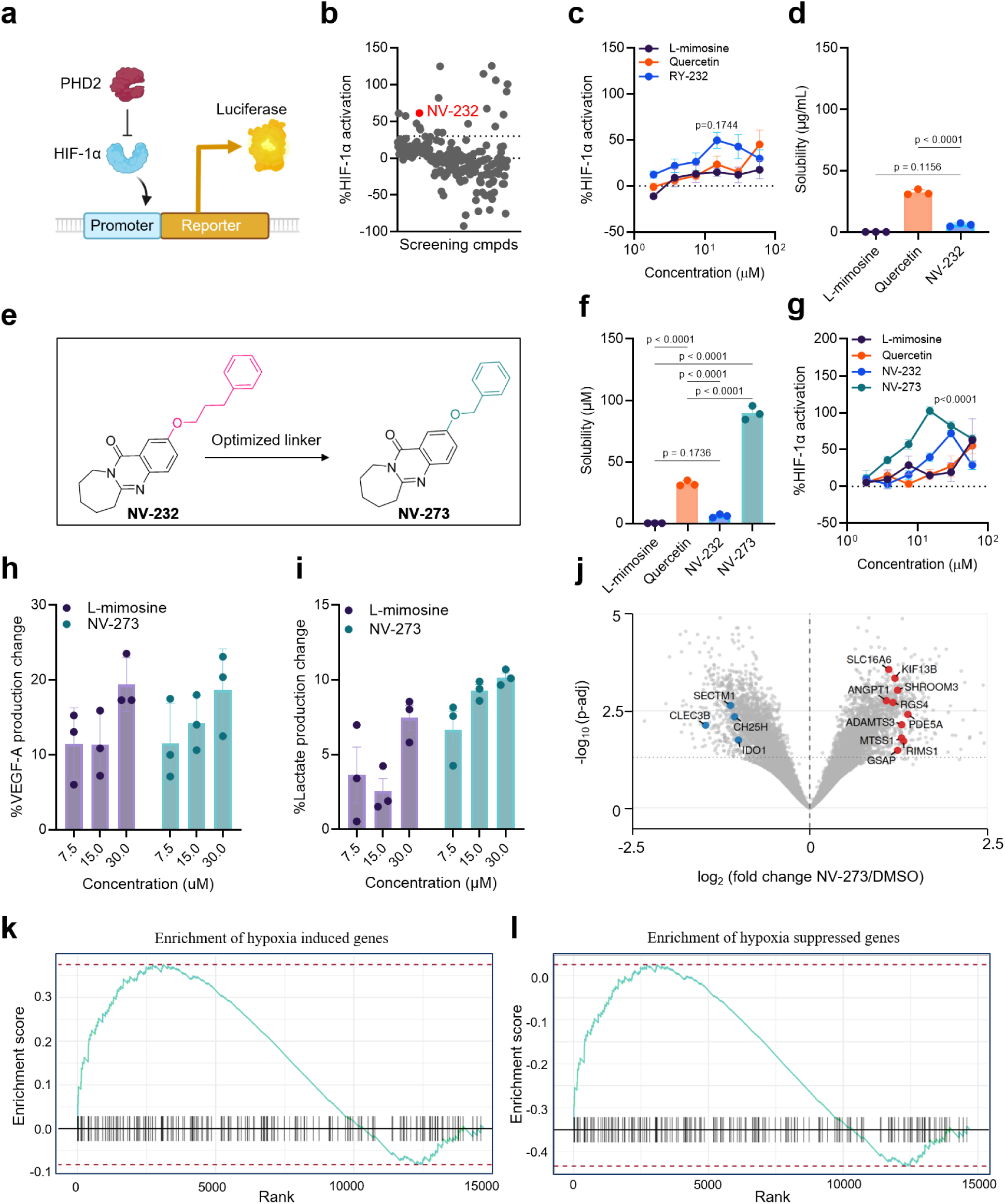
NV-273 activated HIF-1α with broad downstream transcriptional responses via PHD2 inhibition. (a) Principle of the HRE-Luc reporter assay. (b) Change in HIF-1α activation of virtual screening-prioritized compounds in HRE-Luc reporter cells. (c) The top hit NV-232 showed stronger PHD2 inhibition than the benchmark compound L-mimosine after 10 h treatment of HRE-Luc reporter cells. (d) NV-232 is less soluble than the benchmark compound quercetin. (e) Hit-to-lead optimization to improve the aqueous solubility of NV-232 produced NV-273, which features a shorter linker. (f) NV-273 showed higher aqueous solubility than NV-232 and quercetin. (g) NV-273 demonstrated greater HIF-1α activation than the parent compound NV-232 and L-mimosine after 24 h of treatment. NV-273 promoted (h) VEGF-A production in HEKs and (i) lactate production in DPCs after 24 h treatment with higher lactate production than L-mimosine. (j) RNA sequencing revealed increased expression of HIF-1α target genes in NV-273-treated DPCs, consistent with PHD2 pathway engagement. GSEA showed (k) positive enrichment of the manalo_hypoxia_up gene set in NV-273–treated cells relative to DMSO controls (NES = 1.5, p_adj_ = 7.9 × 10⁻⁴) and (l) negative enrichment of the manalo_hypoxia_dn gene set (NES = −1.8, p_adj_ = 7.8 × 10⁻⁹), indicating activation of hypoxia-responsive programs. Data in (a) are representative of the mean of n=3 technical replicates from >3 experimental replicates. Data in (c-d and f-i) represent mean ± s.d. from n=3 technical replicates and are representative of >3 experimental replicates. Data in (h) are derived from HEK donor 1. Data in (i) are derived from DPC donor 1. Data in (j–l) are representative of two experimental replicates from donor 1 and donor 2. p values for pairwise comparisons between indicated groups are shown above bars in (d, f, h, and i), calculated using one-way ANOVA followed by Tukey’s multiple comparisons test and above 15 µM point (c and g), calculated using ordinary two-way ANOVA.

Because L-mimosine is not widely used in consumer products, further characterization and validation were also benchmarked against quercetin, a commonly used product ingredient with PHD2 inhibitory activity^40^. NV-232 emerged as the most efficacious initial hit, producing 50% maximal HIF-1α activation after 24 h of treatment (Fig. 3c). In contrast, L-mimosine elicited slightly weaker activation (p = 0.1744) at 15 µM, possibly due to its poor cell permeability^41^. However, solubility measurements showed that NV-232 was poorly soluble in water (6.1 μM solubility) – comparable to L-mimosine (0.22 μM, p = 0.1736) but lower than quercetin (32 μM, p = 0.0002; Fig. 3d). To overcome this limitation, we performed a computational hit-to-lead optimization to improve the hydrophilicity of NV-232 through computational catalog expansion and obtained NV-273, which has a shortened linker between the terminal aromatic ring and the core tricyclic scaffold (Fig. 3e). NV-273 displayed a 15-fold improvement in aqueous solubility (90 μM) relative to NV-232 and was significantly more soluble than quercetin (p < 0.0001; Fig. 3f). Notably, NV-273 was soluble at 40,000 μM in ethoxydiglycol, a common solubilizer^42^, suggesting that NV-273’s aqueous solubility may be further improved through rational formulation (Supplementary Fig. 3b).

NV-273 exhibited superior PHD2 inhibitory capacity, inducing a 102% increase in HIF-1α activation at 60 μM after 24 h of treatment, exceeding both NV-232 and L-mimosine at 15 µM (p < 0.0001; Fig. 3g). Importantly, the activity of NV-273 was faster and more sustained, with significant HIF-1α stabilization observed at 10 h and maintained up to 48 h post-treatment (Supplementary Fig. 3c), indicating prolonged PHD2 inhibition and subsequent HIF-1α activation. In contrast, NV-232 and L-mimosine showed attenuated or transient activation by 24 h (Supplementary Fig. 3c).

We observed that NV-273 did not induce detectable primary macrophage activation, genotoxicity, skin sensitization, or ROS production (Supplementary Figs. 4a-d). Quercetin, on the other hand, led to pronounced ARE-Nrf2 activation and elevated ROS levels, suggesting a higher sensitization potential, consistent with previous findings^43^. Neither NV-273 nor quercetin substantially affected the viability of most primary cells (Supplementary Fig. 4e, f); however, quercetin modestly increased HDF proliferation at high concentrations and PBMC metabolic activity at low concentrations.

Lactate production from DPCs and VEGF-A production from HEKs were measured as functional readouts of HIF-1α-associated activity, capturing mesenchymal metabolic remodeling and epithelial angiogenic signaling, respectively. Consistent with robust HIF-1α activation, treatment of primary human DPCs and HEKs with NV-273 resulted in a dose-dependent increase of extracellular lactate and VEGF-A production, respectively. Specifically, NV-273 increased VEGF-A production in primary HEKs by 18.7% and elevated extracellular lactate production in primary DPCs by 10.2%, relative to vehicle-treated controls (Figs. 3h and i). These results further supported PHD2 pathway engagement of NV-273 and the capacity of NV-273 to promote downstream regenerative programs consistent with HIF-1α activation.

To further confirm these findings, we performed RNA-sequencing following NV-273 treatment and compared differential gene expression (DEG) relative to vehicle-treated controls (0.1% DMSO). Gene set enrichment analysis (GSEA) revealed that NV-273 and L-mimosine induced similar gene up- and down-regulation consistent with HIF-1α activation (Supplementary Fig. 5)^44^. The sequencing results showed 809 significant DEGs. Because HIF-1α activation has been reported to drive broad transcriptional remodeling, the large number of DEGs observed here is expected. In addition, we identified a subset of annotated genes whose regulation was consistent with PHD2 inhibition (Fig. 3j). Among the upregulated genes, ANGPT1 (angiopoietin 1), ADAMTS3 (ADAM metallopeptidase with thrombospondin type 1 motif 3), RGS4 (regulator of G protein signaling 4), and SLC16A6 (solute carrier family 16 member 6) are associated with hypoxia-adaptive responses, extracellular remodeling, and vascular-supportive programs that are broadly consistent with HIF-1α stabilization and its downstream signaling programs. The induction of ANGPT1 is consistent with activation of a pro-angiogenic tissue-responsive state that complements canonical HIF–VEGF axis signaling. Additional induced genes, including SHROOM3 (shroom family member 3), KIF13B (kinesin family member 13B), PDE5A (phosphodiesterase 5A), MTSS1 (MTSS I-BAR domain containing 1), RIMS1 (regulating synaptic membrane exocytosis 1), and GSAP (gamma-secretase activating protein), further indicate broad transcriptional remodeling after treatment. In parallel, downregulation of IDO1 (indoleamine 2,3-dioxygenase 1), CH25H (cholesterol 25-hydroxylase), SECTM1 (secreted and transmembrane 1), and CLEC3B (C-type lectin domain family 3 member B) suggests suppression of a subset of immune- and signaling-related transcripts. GSEA further supported activation of hypoxia-associated transcriptional programs, with significant enrichment (NES = 1.5, p_adj_ = 7.9 × 10⁻⁴) and reciprocal negative enrichment (NES = −1.8, p_adj_ = 7.8 × 10⁻⁹) of the Manalo hypoxia databases^45^, consistent with coordinated induction of hypoxia-responsive genes and suppression of transcripts normally reduced under hypoxic conditions (Figs. 3k, l). Collectively, these results establish NV-273 as a potent and developable PHD2 inhibitor that drives consistent HIF-1α activation and promotes favorable pro-angiogenic and metabolic responses in primary cells.

### NV-1065 is a non-steroidal 5α-reductase inhibitor and rescued testosterone-induced hair follicle growth inhibition

To identify non-steroidal 5-AR inhibitors, we established a competitive ELISA that quantifies residual testosterone following enzymatic conversion, allowing a direct measurement of 5-AR enzyme activity (Fig. 4a).

**Figure 4.**
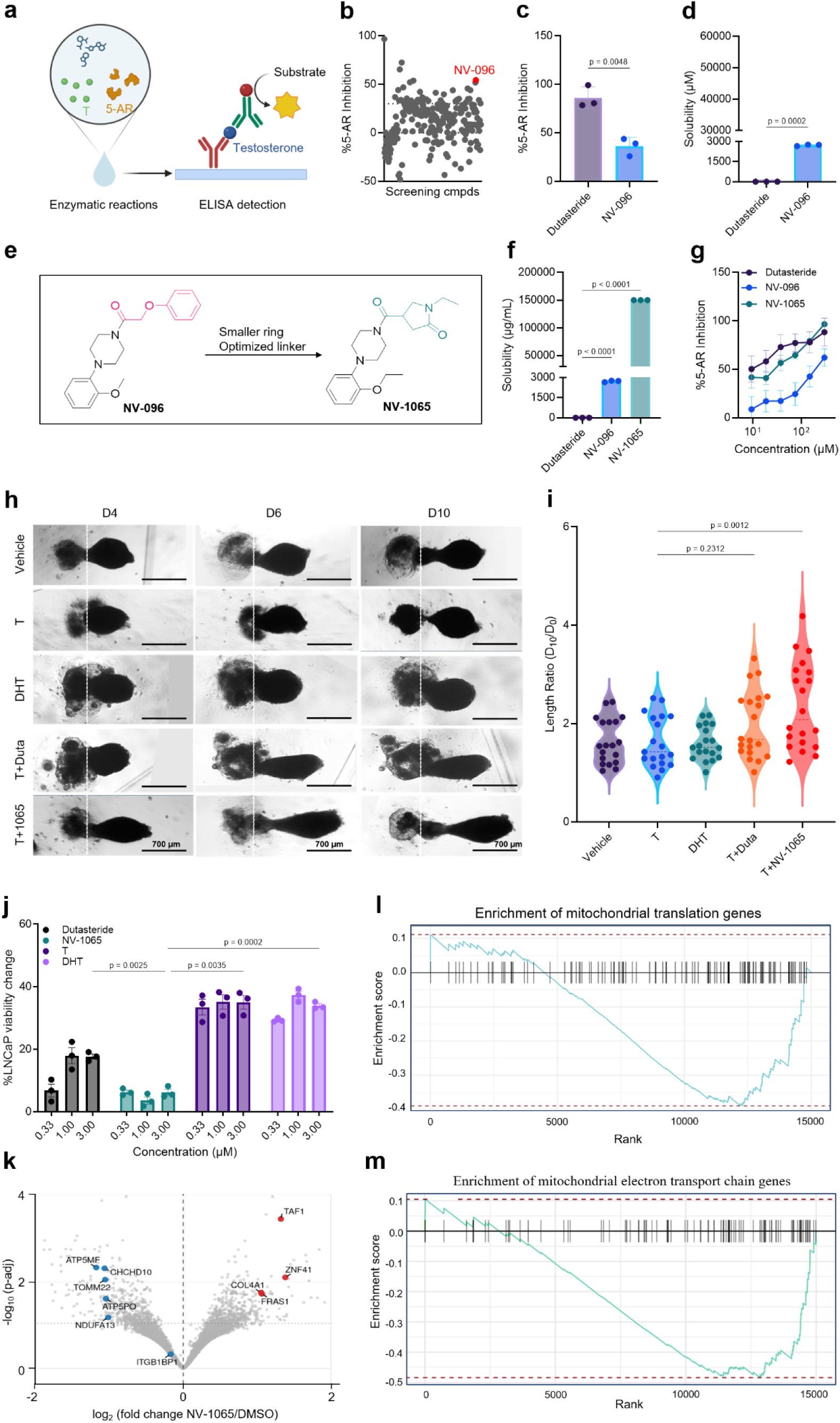
Discovery, optimization, and characterization of non-steroidal 5-AR inhibitors. (a) Schematic of the competitive ELISA used for 5-AR inhibitor screening. Residual testosterone from completed enzymatic reactions competes with testosterone-HRP for binding to a limiting detection antibody. (b) Summary of 5-AR inhibition for compounds prioritized through virtual screening. (c) Inhibitory activity of the top hit NV-096 compared with the benchmark compound dutasteride at 20 µM. (d) NV-096 is more water soluble than dutasteride. (e) Hit optimization to improve NV-096 aqueous solubility. (f) NV-1065 exhibited greater aqueous solubility than NV-096. (g) NV-1065 demonstrated improved 5-AR inhibition relative to the parent compound NV-096 and achieved comparable potency to the benchmark compound dutasteride. (h) Representative images and (i) quantitative analysis showing NV-1065 rescued testosterone-induced growth inhibition in human follicle organoids.The white dashed line marks the position from which the follicle organoid sprouting initiates. T: testosterone; DHT: dihydrotestosterone; Duta: dutasteride. (j) NV-1065 did not stimulate proliferation of the steroid-sensitive prostate cancer cell line LNCaP, whereas testosterone, DHT, and dutasteride induced significant growth. (k) RNA sequencing revealed that NV-1065 reduced AR-mediated transcription in testosterone-treated DPCs. GSEA of (l) reactome_mitochondrial_translation (NES = -1.5, p_adj_ = 5.3 × 10⁻³) gene set showing reduced mitochondrial protein synthesis and (m) wp_electron_transport_chain_oxphos_system_in_mitochondria (NES = −1.8, p_adj_ = 5.8 × 10⁻⁵) gene sets showing suppression of oxidative phosphorylation, which are consistent with reduced AR-mediated signaling and transcription. Data in (b) are representative of the mean of n=3 technical replicates from >3 experimental replicates. Data in (c-d), (f-g) and (i) represent mean ± s.d. from n=3 technical replicates. Data in (i) represent mean ± s.d. from n=19 (vehicle, testosterone, and DHT) or n=20 (T + dutasteride and T + NV-1065) technical replicates. Data in (k-m) are derived from two experimental replicates of n=2 biological replicates. p values for pairwise comparisons between indicated groups are shown above bars, calculated using one-way ANOVA followed by Tukey’s multiple comparisons test.

The 5-AR inhibitory activity of the virtual screening-prioritized compounds was evaluated using human liver S9 fractions containing active 5-ARs. Percentage inhibition was determined by normalizing compound-treated enzyme reactions to vehicle-treated (0.1% DMSO) reactions (Fig. 4b). As a starting point, compounds exhibiting greater than 30% inhibition at 30 μM were prioritized. NV-096 emerged as an initial hit, achieving 36% inhibition of 5-ARs at 20 μM whereas dutasteride achieved 79% inhibition (Fig. 4c). Although less potent than dutasteride, NV-096 harbors no steroidal backbone and therefore provides a non-steroidal structural starting point for further optimization. LC-MS studies revealed that NV-096 exhibited limited aqueous solubility (2,717 μM), albeit higher than dutasteride (3.6 μM) (p = 0.0002, Fig. 4d).

The computational hit-to-lead optimization process focused on modifying the terminal aromatic ring and ester linker to increase hydrophilicity while preserving 5-AR inhibitory activity (Fig. 4e). This effort yielded the optimized analogue NV-1065, which displayed a significant enhancement in aqueous solubility, exceeding 150,000 μM and surpassing both the parent compound NV-096 and dutasteride by several orders of magnitude (p < 0.0001; Fig. 4f). Meanwhile, NV-1065 exhibited substantially improved 5-AR inhibition relative to NV-096 and achieved nearly full inhibition at high doses, comparable to dutasteride (p = 0.8755; Fig. 4g), indicating that improvements in solubility did not compromise its 5-AR inhibitory capacity. Additionally, we evaluated the mode of inhibition of NV-1065 by comparing its potency with or without 5-AR preincubation (Supplementary Fig. 6a). As expected, dutasteride exhibited *irreversible* inhibition as evidenced by improved potency with 5-AR preincubation (Supplementary Fig. 6b). Negligible changes in potency were observed for NV-1065 regardless of 5-AR preincubation, suggesting a *reversible* inhibition mechanism.

Next, we investigated whether NV-1065 could mitigate androgen-induced growth suppression in human hair follicle organoids. As DPCs and HEKs express 5-ARs^46,47^, exogenous testosterone can be converted into DHT, thereby limiting organoid elongation. Over a 10-day culture period, treatment with testosterone or DHT revealed a trend toward reduced organoid elongation relative to vehicle controls, consistent with their role in hair follicle miniaturization (Fig. 4h). Under the same conditions, dutasteride treatment partially rescued testosterone-induced growth inhibition (p>0.2312). In contrast, the addition of NV-1065 in testosterone-treated organoids markedly restored elongation compared with testosterone alone (p = 0.0012) (Fig. 4h and i).

Because NV-1065 lacks a steroidal backbone, we next assessed whether it showed steroid-like cellular activity, using the androgen-sensitive prostate cancer cell line lymph node carcinoma of the prostate (LNCaP), in which cell proliferation can be stimulated by steroids such as testosterone and DHT^48^. Consistent with its non-steroidal scaffold, NV-1065 did not promote LNCaP cell proliferation, whereas testosterone, DHT and dutasteride promoted LNCaP cell proliferation by up to 35%, 37%, and 18%, respectively (Fig. 4j), significantly greater than that induced by NV-1065 (p < 0.005). These data highlight NV-1065 as a non-steroidal 5-AR inhibitor. Moreover, NV-1065 exhibited a favorable cellular specificity profile across multiple primary cell types. Comparable to dutasteride, NV-1065 did not induce detectable macrophage activation, genotoxicity, skin sensitization, or intracellular ROS accumulation (Supplementary Figs. 7a-d). Furthermore, NV-1065 did not produce significant changes in the CTG or CTB response of various primary skin cells or non-skin primary cells (Supplementary Figs. 7e,f). In contrast, dutasteride elicited pronounced cytotoxicity in HEKs at 30 μM, highlighting a differential tolerability profile at higher concentrations (Supplementary Fig. 7e).

RNA sequencing identified a set of 144 DEGs in NV-1065-treated cells relative to vehicle treatment, including a substantial subset overlapping an SRD5A1/2-consistent expression signature. This transcriptional shift is consistent with suppression of DHT-dependent dermal papilla signaling. Moreover, treatment with NV-1065 resulted in a pathway enrichment similar to that observed with dutasteride, with perturbation of mitochondrial and cholesterol metabolic pathways consistent with hallmark 5-AR inhibition (Supplementary Fig. 8). Among the most notable changes were reduced expression of genes involved in transcriptional regulation and extracellular niche organization, including TAF1 (TATA-box binding protein-associated factor 1), ZNF41 (zinc finger protein 41), COL4A1 (collagen type IV alpha 1 chain), FRAS1 (Fraser extracellular matrix complex subunit 1), and ITGB1BP1 (integrin beta-1-binding protein 1), in line with prior evidence that androgen signaling reshapes dermal papilla gene expression and paracrine output. Additionally, a cluster of mitochondrial OXPHOS components, ATP5MF (ATP synthase membrane subunit f), ATP5PO (ATP synthase peripheral stalk subunit OSCP), CHCHD10 (coiled-coil-helix-coiled-coil-helix domain-containing protein 10), NDUFA13 (NADH:ubiquinone oxidoreductase subunit A13), and TOMM22 (translocase of outer mitochondrial membrane 22) were significantly downregulated, in line with the known dependence of mitochondrial steroidogenesis on membrane integrity and electron transport chain function. GSEA further indicated suppression of mitochondrial metabolic programs following NV-1065 treatment. Both the mitochondrial protein synthesis (NES = −1.5, p_adj_ = 5.3 × 10⁻³) and mitochondrial oxidative phosphorylation (NES = −1.8, p_adj_ = 5.8 × 10⁻⁵) gene sets showed negative enrichment. This pattern is consistent with a coordinated downregulation of energy-intensive biosynthetic and respiratory pathways, which are often supported by androgen signaling. Collectively, these results establish NV-1065 as a potent, highly soluble non-steroidal 5-AR inhibitor that lacks detectable androgenic activity in the tested cellular assays and restores growth in androgen-suppressed follicle organoids.

### Combinatorial efficacy of lead compounds in hair follicle rejuvenation

To determine whether the four lead compounds could be combined without compromising solubility, compound compatibility, or multi-pathway activity, we developed a water-based solution containing 0.10% each of NV-623, NV-624, and NV-1065, and 0.05% of NV-273. The solvents included ethanol, dimethyl isosorbide, polysorbate 20, 1,3-propanediol, ethoxydiglycol in water (w/w, Fig. 5a). This aqueous format was selected to enable simultaneous incorporation of all four compounds and direct evaluation of their combined *in vitro* efficacy while preserving the chemical integrity of each component. Long-term stability under ambient storage was monitored by periodic LC–MS quantification over 150 days, during which all four compounds showed minimal degradation, with more than 90% of the initial amount remaining (Fig. 5b).

**Figure 5.**
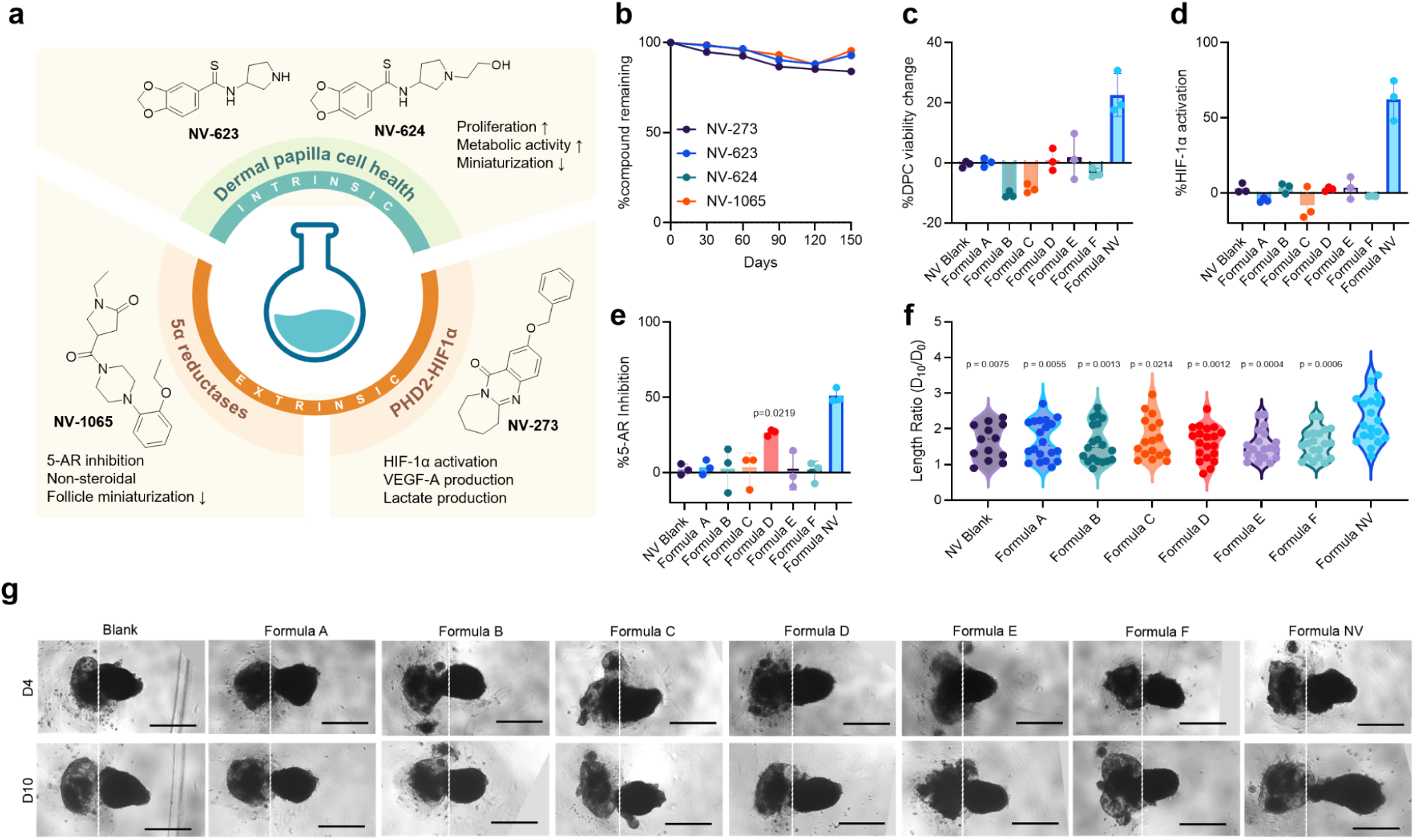
Combinatorial efficacy of top leads in hair follicle rejuvenation. (a) Scheme of lead compound formulation. NV-623, NV-624, and NV-1065 were added at 0.10% (w/w), and NV-273 was added at 0.05%. The formulation was optimized to maximize the solubility and stability of all four compounds. (b) All four lead compounds showed high stability. The formula was left under ambient conditions for over 90 days and showed >90% remaining after 150 days. (c) The formula NV exhibited strong efficacy in promoting DPC proliferation. (d) The formula NV exhibited strong efficacy in PHD2 inhibition. (e) The formula NV exhibited strong 5-AR inhibitory effects. (f) Quantitative analysis and (g) representative images showing formula NV promoted follicle organoid elongation. The white dashed line marks the position from which the follicle organoid sprouting initiates. All formulas were tested at 500-fold dilution. Data in (c) represent mean ± s.d. from 3 experimental replicates of 3 independent donors. Data in (d-e) represent mean ± s.d. of n=3 technical replicates from 3 experimental replicates. Data in (f) represent n = 13 (NV Blank), n = 20 (formulas A, D and F), n = 19 (formulas B, E, and formula NV), n = 17 (formula C) from over 3 experimental replicates. Images in (g) are representative of over 3 experimental replicates of 10 technical replicates. p values are indicated above bars for comparisons to the other formulas, calculated using one-way ANOVA with Tukey’s test. Formula A was obtained from Kérastase; formula B was obtained from Equate^TM^; formula C was obtained from Nutrafol; formula D was obtained from The Ordinary; formula E was obtained from Vegamour; formula F was obtained from KilgourMD.

To confirm that compound efficacy was retained after formulation, we benchmarked the formula (formula NV) against commercially available hair thinning and hair loss products spanning distinct active ingredients including minoxidil, caffeine, botanical-based extracts, peptides, and plant exosomes. We identified a concentration with minimal solvent interference (Supplementary Figs. 9a-c), and tested the functional activity of the formula NV using DPCs, HRE-Luc reporter cells, and human liver S9 fractions containing active 5-AR enzymes. At a 1600-fold dilution, formula NV increased DPC viability by 23% relative to vehicle (water-treated) control (p<0.0001 relative to other formulas; Fig. 5c). Among the other commercial formulas, only formula A showed a modest 12% increase in the CTG response in DPCs. Formulas D and E exhibited negligible changes while formulas B and C reduced DPC viability by around 6.0% under the same conditions. Furthermore, formula NV induced robust HIF-1α stabilization, reaching 97% HIF-1α activation at the same dilution relative to vehicle control, indicating effective inhibition of PHD2 activity (p < 0.0001 relative to other formulas; Fig. 5d). Meanwhile, formulas B and D showed around 15% HIF-1α activation while formulas C and E showed no detectable effects. In the 5-AR assay, formula NV inhibited 5-AR enzyme activity by 74% at a 1600-fold dilution relative to vehicle control, whereas the other formulas showed less than 40% inhibition (Fig. 5e, p = 0.0219 relative to formula D; p < 0.0001 relative to other formulas). The 5-AR inhibition from the commercially available products may also be due to the various ingredients’ capacity to inactivate or denature proteins from the liver fractions.

The functional efficacy of formula NV was further evaluated in the follicle organoid model and benchmarked against the same set of commercial formulas and individually formulated compounds. Consistent with the activity of the non-formulated compounds (Figs. 2–4), the individually formulated compounds retained their expected efficacy (Supplementary Figs. 9d, e). Formula NV, which contains all four lead compounds, further significantly increased follicle organoid sprouting length compared with blank (formula solvents only) treatment (Figs. 5f, g; p = 0.0075). This effect was also greater than that of all commercial formulas (p < 0.03 for all comparisons), which showed no measurable organoid elongation under the same conditions. Together, these results demonstrate that the integrated formula NV preserved the combinatorial activities of the lead compounds across DPC proliferation, PHD2 inhibition, 5α-reductase inhibition, and promotion of follicle organoid sprouting.

## Discussion

Most interventions for androgenetic alopecia are suboptimal or have undesirable side effects. Here, we show how an AI-enabled discovery framework can help address this gap by uncovering compounds that act across three complementary pathways relevant to hair follicle rejuvenation. Using this framework, we identified hits and optimized lead compounds that support DPC viability, activate hypoxia-responsive regenerative signaling, and suppress androgen-mediated follicular miniaturization through 5α-reductase inhibition.

Unlike prednisolone, which broadly potentiates primary cell proliferation and suppresses immune responses, the optimized DPC-active compounds NV-623 and NV-624 combined robust activity with low promiscuity and low cytotoxicity. Both compounds also significantly promoted follicular elongation in a hair organoid model, with clear differences observable within 10 days. NV-273 produced stronger and more sustained HIF-1α activation and increased secretion of VEGF-A and lactate than L-mimosine and quercetin. For androgen-pathway modulation, the non-steroidal compound NV-1065 inhibited 5α-reductase activity and rescued androgen-induced growth suppression in the follicle organoid model. RNA-seq analysis provided orthogonal support for target engagement at the transcriptional level. Treatment with the PHD2 inhibitor NV-273 induced genes associated with vascular support, extracellular remodeling, and metabolic adaptation, including ANGPT1, which is linked to Tie-mediated angiogenic signaling; ADAMTS3, an extracellular metalloprotease associated with matrix organization and collagen processing; and SLC16A6, a plasma-membrane monocarboxylate transporter involved in organic acid transport^49–51^. In parallel, NV-1065 treatment was associated with suppression of androgen-responsive transcriptional programs such as blunted DHT-driven inhibitory mediators DKK1 (Dickkopf WNT signaling pathway inhibitor 1) and TGF-β (transforming growth factor-β)–associated signals in androgen-responsive follicular cells.

Several limitations should be noted. Although RNA-seq supported target engagement for NV-273 and NV-1065, some of the most pronounced transcriptional changes extended beyond the intended pathways, reflecting broader cellular responses to treatment. For example, prior toxicogenomic studies demonstrated that compound-induced transcriptomic responses were often enriched for genes involved in xenobiotic handling, including drug metabolism, transport, and broader detoxification-associated stress programs^52,53^. In addition, NV-273 appeared to downregulate several immune-related genes and both dutasteride and NV-1065 showed enrichment of androgen-response signatures in the 5α-reductase RNA-seq datasets. This could reflect effects on other pathways besides the bias towards the pathways of interests. Interpretation is also limited by the use of 2D cell cultures for RNA-seq, where pathway-level trends were evident even when individual changes were modest^44,54^. Additionally, because the organoid model cannot wholly represent the physiological environment, NV-273 cannot be thoroughly assessed, and NV-1065 can only be assessed with exogenous testosterone, limiting physiological relevance. Further mechanistic studies will be needed to distinguish direct from adaptive effects and to define the temporal consequences of 5α-reductase modulation in more physiologically relevant systems. Future studies should assess long-term efficacy and safety in clinical settings to further support the translational potential of the lead compounds. In parallel, clinical assessment of skin tolerability and sensitization risk will be important to establish their tolerability. Expanded transcriptomic and proteomic profiling will also be important for refining mechanistic understanding and identifying biomarkers of response, especially for NV-623 and NV-624.

As with other applications, the productivity of AI-enabled discovery approaches relies substantially on the quality of the training data used to develop the models. Here, we have generated custom, experimentally-derived high-throughput screening data at scale to train such models. By integrating phenotypic screening, target-based prioritization, and experimental validation within a single experimental framework, we show how computationally guided discovery can be coupled to biological and translational endpoints to accelerate the discovery of compounds with complementary mechanisms of action. In this context, the value of the approach lies not only in improving hit selection, but also in enabling the systematic assembly of functionally aligned molecules that together address distinct but convergent drivers of tissue decline. More broadly, this work provides a generalizable framework for the discovery of mechanistically integrated bioactive compounds and supports the broader application of AI-enabled, multi-target design strategies to other complex biological systems.

## Materials and methods

### Cells

Human dermal papilla cells (DPCs; Cell Applications, 602K-05a), human dermal fibroblasts (HDFs; Cell Applications, 106K-05a), human dermal microvascular endothelial cells (HDMECs; PromoCell, C-12210), human epidermal keratinocytes (HEKs; Cell Applications, 102-05a), human preadipocytes (HPADs; Cell Applications, 802s-05a), human renal cortical epithelial cells (HRCEpCs; PromoCell, 501Z019.23), and human skeletal muscle cells (HSKMCs; Cell Applications, 150K-05a) were obtained from the indicated commercial sources and cultured according to the manufacturers’ recommendations. Peripheral blood mononuclear cells (PBMCs) were purchased from HumanCells Biosciences (PBMC-C100M) or obtained from de-identified leukopaks (BioIVT, HUMANLMX25-0001127) following enrichment in a Ficoll gradient. All available donor information is listed in **Supplementary Table 1**. To generate the hypoxia-response reporter cells, we stably integrated a hypoxia response element (HRE)–driven luciferase reporter via lentiviral transduction (BPS Bioscience, 78668) into HEK-293T cells (ATCC, CRL-3216). ARE-Nrf2 HepG2 reporter cells (BPS Bioscience, 60513) were used to evaluate Nrf2 pathway activation and maintained based on the manufacturers’ recommended protocols.

### Reagents

Follicle dermal papilla cell growth medium (PromoCell, C-26501), renal epithelial cell growth medium (PromoCell, C-26130), human skeletal muscle cell medium (Cell Applications, 151-500), human EpiVita medium (Cell Applications, 141-500a), human preadipocyte medium (Cell Applications, 811-500), human CADMEC medium (Cell Applications, 112-500), and human dermal fibroblast medium (Cell Applications, 116-500) were used as specified. Basal media included DMEM-GlutaMAX™ (Thermo Fisher Scientific, 10569010), Advanced DMEM/F-12 (Thermo Fisher Scientific, 12634010), and RPMI-1640 (Thermo Fisher Scientific, 11875093). Additional reagents were purchased from Thermo Fisher Scientific: Fetal bovine serum (FBS; A5670801), penicillin–streptomycin (15140122), Trypsin-EDTA (25200056), TrypLE™ Express (12604021), and DPBS (14190250). 10× PBS was obtained from Alkali Scientific (PB174). Trypan blue solution (25-900-CI) and Matrigel (356255) were obtained from Corning.

CellTiter-Blue® cell viability assay (CTB; Promega, G8081), CellTiter-Glo® 2.0 cell viability assay (CTG; Promega, G9243), ROS-Glo™ H_2_O_2_ assays (Promega, G8821), Bright-Glo™ luciferase assay system (Promega, E2650), and Lactate-Glo™ assays (Promega, J5022) were performed according to the manufacturers’ protocols. Testosterone concentrations were measured using a testosterone ELISA kit (Arbor Assays, K032-H5), and VEGF-A levels were quantified using a human VEGF-A DuoSet ELISA kit (Bio-Techne, DY293B-05). The Promega™ GloMax® Plate Reader was used for the luminescence, fluorescence, and absorbance measurements. Countstar Castor X1 high-throughput imager was used for DAPI imaging.

Human Leukopak (HUMANLMX25-0001127) was purchased from BioIVT for PBMC enrichment. CD14^+^ monocytes were further isolated using the human pan monocyte isolation kit (480059) from Biolegend. Immuno-SF macrophage media (10961) from StemCell Technologies were supplemented with recombinant human M-CSF protein (Sino Biological, 11792-HNAH-20) for monocyte differentiation into macrophages. Lipopolysaccharide (LPS; 00-4976-93) from Thermo Fisher was used to activate the unpolarized macrophages. Zombie violet (423113), CD14-AF488 (301817), HLA-DR-PE (307605) from Biolegend were used to stain macrophages for flow cytometry.

NV-168 was purchased from Molport. NV-232 and NV-273 were purchased from Vitas. NV-1065 was purchased from ChemBridge. 6-bromoindirubin-3′-oxime (Millipore Sigma, B1686), recombinant human basic fibroblast growth factor (bFGF; Sino Biological, 10014-HNAE-100), and recombinant human bone morphogenetic protein-2 (BMP2; Sino Biological, 10426-HNAE-100) were used to preserve physiological signaling in adult dermal papilla cells^55^. 4′,6-Diamidino-2-phenylindole (DAPI; Biotium, 40011) was used for nuclear staining. Paraformaldehyde (PFA; MP Biomedicals, 0219998320) was used for cell fixation. Bovine serum albumin (BSA; GoldBio, A-420-10) was used to block non-specific protein binding. Human liver S9 fraction was used as a source of 5-alpha reductase and was obtained from Thermo Fisher Scientific (HMS9PL).

FastPure Cell/Tissue Total RNA Isolation Kit V2 (Vazyme, RC-112-01) was used for RNA extraction from cell cultures. Tween-20 was purchased from Dot Scientific (DSP20370-0.5), and Triton X-100 (648466-50ML) was obtained from Millipore Sigma. Sodium hydroxide (NaOH; Spectrum Chemical, S1310-100MLPL) was used for pH adjustment. LB broth (VWR, 89500-596) and carbenicillin (VWR, J358-1G) were used for bacterial culture where indicated. 4-Nitroquinoline N-oxide (SigmaAldrich, N8141-250MG), 5-bromo-4-chloro-3-indolyl-β-D-galactopyranoside (SigmaAldrich, 9660-1GM), minoxidil sulfate (Cayman Chemical, 36084-10), prednisolone disodium acetate (Chemscene, CS-2878_100mg), quercetin dihydrate (AstaTech, 41708-25G), L-mimosine (Cayman Chemical, 14337-50), NADPH tetrasodium (Ambeed, A341469-250mg), testosterone (Cayman Chemical, ISO60154), dihydrotestosterone (DHT; Cayman Chemical, 41649), dutasteride (MedChemExpress, HY-13613), and finasteride (Cayman Chemical, 14938-50) were obtained from the indicated vendors and prepared according to the manufacturers’ specifications.

Formula A (Genesis Serum Fortifiant Hair Serum) was obtained from Kerastase. Formula B (Hair Regrowth Treatment for Women, 2% Minoxidil Topical Solution) was obtained from Equate^TM^. Formula C (Women Hair Serum) was obtained from Nutrafol. Formula D (Multi-Peptide Serum for Hair Density) was obtained from The Ordinary. Formula E (GRO Hair Serum) was obtained from Vegamour. Formula F (The Treatment) was obtained from KilgrourMD.

### AI-enabled discovery framework

Two parallel discovery workflows were used to identify small-molecule modulators of intrinsic and extrinsic pathways for hair follicle rejuvenation. For the cell-intrinsic pathways, we implemented a phenotype-guided, machine learning–assisted small-molecule workflow in primary human DPCs, as described in previous work^23–25^. A curated small-molecule library of 5×10^4^ compounds was assembled to broadly sample chemical space, taking into account structural diversity, physicochemical properties, and known toxicity or undesirable bioactivity. A robotic high-throughput screening (HTS) experiment was performed using the compounds at a final concentration of 10 μM to quantify the changes in cell metabolic activity and proliferation in two DPC donors. CTB and CTG reagents were used as proxies for metabolic activity and proliferation, respectively. The resulting HTS dataset was used to train an ensemble of Chemprop models^56^. After training, the ensemble served as a phenotype-aware predictor of DPC responses (based on CTG/CTB readouts) and was used to computationally prioritize compounds from a larger database of ∼11 million commercially available compounds from MolPort, a chemical provider.

For the cell-extrinsic pathways, we compiled a library of approximately 17 million compounds from public and open-source collections provided by chemical vendors and research institutes. The sequence for PHD2 was obtained from the UniProt entry Q9GZT9 and the sequence for 5AR was obtained from the UniProt entry P18405. We then implemented a two-step virtual screening workflow that incorporated multiple screening methods at each stage to preserve chemical diversity. This strategy helped reduce target- and method-specific biases that may arise when relying on a single screening approach. In the primary screen, compounds were filtered by physicochemical criteria (250 < MW < 450; logP < 3.5) and then evaluated using three deep learning drug–target affinity (DTA) models. ∼10,000 candidates were prioritized from the list using an even-weighted selection strategy to maintain chemical diversity across scoring distributions.

Secondary screening was performed using multiple methods in parallel, including the machine-learning model DiffDock and the physics-based MM/PBSA method^57–59^. The curated candidate set was screened concurrently alongside a panel of known target inhibitors included as positive controls. Compounds were prioritized based on (i) screening scores relative to the positive-control inhibitors and (ii) method-specific confidence criteria.

### Computational hit-to-lead optimization and medicinal chemistry

For select targets, computational hit-to-lead optimization was performed using two parallel strategies: (1) similarity-based searching against in-house molecular libraries and (2) pharmacophore-informed molecular generation with TransPharmer^60^. In the similarity-based approach, RDKit fingerprints and Tanimoto similarity coefficients were used to identify derivative compounds. A similarity threshold of 0.8 was used. In parallel, TransPharmer was applied using the generate_pc.yaml configuration file according to the authors’ recommendation to generate candidate compounds from pharmacophore fingerprints. All generated candidates were then filtered to retain compounds with molecular weight < 450 g/mol and logP < 3.5.

Medicinal chemistry of NV-168 was performed by coupling hydrophilic building blocks or tethering linkers of different structures to the thioamide core. The thioamide core scaffold was prepared using Lawesson’s reagent with the amide core^61^. LC-MS and NMR characterization of NV-623 and NV-624 can be found in the Supplementary Spectra.

### In vitro assays

For DPC-based assays, primary human DPCs were plated in complete follicle dermal papilla cell growth medium (PromoCell) supplemented with 1% penicillin-streptomycin (Thermo Fisher), 200 ng/mL BMP-2, 20 ng/mL bFGF, and 1 µM BIO at a density of 700-850 cells per 25 μL/well of 384-well plates and allowed to attach for 12-14 h. The cultures were then treated with various test compounds for at least 36 h. Cell viability was measured following the addition of CTB or CTG as a proxy for cell viability, according to the manufacturer’s protocols. For cell counts, culture media were carefully aspirated, and the cells were washed with 20 μL PBS twice followed by fixation using 10 μL of 1% PFA (MP Biomedicals) at room temperature for 10 minutes. The PFA solution was aspirated, and the cells were washed with 20 μL PBS twice followed by incubation with 10 μL of DAPI (1:5000, Biotium) at room temperature for 15 minutes. Cells were imaged using a Countstar Castor X1 imager.

For the steroidal property test, LNCaP cells were plated in RPMI-1640 media supplemented with 1% penicillin-streptomycin at a density of 5,000 cells per 25 μL/well of 384-well plates and allowed to attach for 24 h. The cultures were treated with test compounds for at least 48 h before cell viability was quantified using CTG.

To evaluate HIF-1α activation via PHD2 inhibition, HRE-Luc-HEK293T reporter cells were plated at a density of 5,000 cells per 25 μL/well of 384-well plates and allowed to attach for 12-16 h. The cells were then treated with the test compounds at various concentrations for 10 h, 24 h, and 48 h. HIF-1α activation was measured using the Bright-Glo^TM^ assay system.

For the 5α-reductase inhibition assay, human liver S9 fractions were diluted to a final protein concentration of 1 mg/mL in PBS buffer (pH 6.5). The diluted fractions were mixed with NADPH (0.2 mM), testosterone (1 µM), and the test compounds to a final volume of 30 µL in round-bottom 96-well plates. The mixtures were incubated at 37 °C for 30 min to enable enzymatic conversion of testosterone to DHT. Reactions were terminated by the addition of 1 N HCl, and samples were diluted 60-fold for ELISA detection. Residual testosterone levels were quantified using a competitive ELISA kit following the manufacturer’s instructions, and inhibitory activity was determined from OD450 values relative to vehicle-treated reactions relative to vehicle-treated reactions.

### Solubility measurements

Relative solubility was assessed by turbidity measurements at OD_600_ ^62^. Compounds were dissolved in Milli-Q water in 384-well plates at a final concentration of 0.25 mg/mL with 2.5% (v/v) DMSO. The absorbance values were normalized to control wells containing 2.5% (v/v) DMSO only.

Quantitative solubility was determined by liquid chromatography–mass spectrometry (LC–MS) using an Agilent 1260 system equipped with a Poroshell 120 column (C18, 2.7 µm, 2.1 × 50 mm). The mobile phases consisted of Milli-Q water and acetonitrile. The samples were acquired in both positive and negative ionization modes.

Calibration curves were generated by plotting integrated peak areas (area under the curve, AUC) from HPLC chromatograms against compound concentrations ranging from 20 to 600 µM. Saturated aqueous solutions were prepared by adding excess solid compound to Milli-Q water, followed by light sonication for 30 min at room temperature. Suspensions were centrifuged at 17,000 × g for 30 min at room temperature, and the supernatants were injected into the LC–MS system. Compound concentrations in saturated solutions were determined by interpolating the measured AUC values against the corresponding calibration curves.

### Lactate and VEGF quantification

Lactate levels in cell culture media were quantified using Lactate-Glo™ according to the manufacturer’s instructions. Briefly, human hair follicle dermal papilla cells (DPCs) or human epidermal keratinocytes (HEKs) were seeded at a density of 10,000 cells per well in DMEM (Thermo Fisher, supplemented with 10% FBS and 1% penicillin/streptomycin) in flat-bottom 96-well plates and cultured at 37 °C in 5% CO_2_. The cells were allowed to attach overnight and the cultures were treated with the test compounds for 16 h. Following treatment, conditioned media were collected and clarified by brief centrifugation to remove cellular debris. Supernatants were diluted 1:200 in phosphate-buffered saline (PBS) for lactate quantification or 1:1 dilution in PBS for VEGF-A quantification. The diluted samples were processed following the standard Lactate-Glo™ protocol or VEGF ELISA protocol, respectively. Lactate concentrations were calculated based on lactate standard curves. All measurements were normalized to vehicle-treated cultures as indicated in the figure legends.

### Hair follicle organoid model

Hair follicle organoids were generated using a co-culture of DPCs and HEKs^33^. Cells were harvested by trypsinization, collected, and resuspended in advanced DMEM/F-12 (supplemented with 10% FBS, 1% penicillin-streptomycin and 2% v/v Matrigel). For organoid formation, 100 µL of the DPC suspension (5.0 × 10³ cells) was first added to each well of round-bottom low-adhesion 96-well plates. Immediately thereafter, 100 µL of the HEK suspension (5.0 × 10³ cells), prepared in the same advanced DMEM/F-12 (10% FBS, 1% penicillin-streptomycin, 2% v/v Matrigel), was gently added to the same wells, resulting in a total of 1.0 × 10⁴ cells per well. Plates were incubated at 37 °C in a humidified 5% CO₂ atmosphere to allow follicle organoid formation. After 2 days, when follicle-like structures were established, compounds were added to each well. A volume of 100 µL of medium was then replaced every 2 days and compounds were replenished in fresh DMEM/F-12 (supplemented with 10% FBS and 1% penicillin-streptomycin). Organoid morphology and length were monitored for up to 10 days using an EVOS microscope (Thermo Fisher), and follicle length was quantified from brightfield images using ImageJ.

### Bulk RNA-sequencing

DPCs were grown in DPC growth medium supplemented with 1% penicillin-streptomycin, 200 ng/mL BMP-2, 20 ng/mL bFGF, and 1 µM BIO at 37 °C in a humidified atmosphere containing 5% CO₂ until ∼75% confluence was reached. The cells were then washed once with PBS, and the medium was replaced with DMEM GlutaMAX supplemented with 10% FBS and 1% penicillin-streptomycin containing 15 µM of the test compounds. Each treatment was performed in two DPC donors (donors 1 and 2; Supplementary Table S1). The DPCs were treated with the test compounds for 12 h. Following treatment, the culture media were carefully aspirated, and cells were immediately lysed *in situ* by adding 0.5 mL lysis buffer from the Vazyme RNA extraction kit per T25 flask. Total RNA was isolated from each sample according to the manufacturer’s instructions. RNA concentration and purity were assessed by NanoDrop prior to downstream library preparation and sequencing. RNA-seq library construction and next-generation sequencing were performed by NGS-lab following standard protocols.

Raw reads were trimmed using Trimmomatic (v0.39) in paired-end mode to remove adapter sequences and low-quality base pairs. Illumina adapters (TruSeq3-PE-2.fa) were removed using the ILLUMINACLIP:2:30:10 setting. Reads were further filtered using LEADING:3, TRAILING:3, and SLIDINGWINDOW:4:15, and reads shorter than 36 bp after trimming were discarded. Trimmed reads were aligned to the human reference genome (GRCh38) using STAR (v2.7.11) with default parameters unless otherwise stated. Alignments were output as coordinate-sorted BAM files, and gene-level read counts were generated using the GeneCounts mode. Gene-level read counts were quantified using featureCounts (Subread v2.1.1) with the GENCODE v49 basic annotation. Reads were counted in paired-end mode, requiring both mates to be mapped (-B). Strand-specific counting was performed in reverse-stranded mode (-s 2). Only primary alignments overlapping exons were assigned to genes based on the gene_id attribute. Differential gene expression analysis was performed using DESeq2 (v1.36) in R. Genes with fewer than 10 counts in at least two samples were removed prior to analysis. DMSO was used as the reference level. Variance-stabilizing transformation (VST) was applied for visualization and clustering. Log2 fold changes were shrunk using the apeglm (v1.16) method. Differentially expressed genes were defined as those with adjusted p-value < 0.05 and |log2 fold change| > 1. Pathway analysis was then performed. The results from differential gene expression analysis were first cleaned by removing rows with missing values and duplicate genes. Over-representation analysis (ORA) was conducted on significantly differentially expressed genes, defined by an adjusted p value < 0.05 when available, or nominal p value < 0.05 otherwise, using the filtered gene set as background. Enrichment was assessed with the upper-tail hypergeometric test, and p values were corrected by the Benjamini–Hochberg method. In parallel, preranked gene set enrichment analysis (GSEA) was performed using a ranked list generated from the signed significance score, *sing*(*log*2 *fold change*) × [− *log*10(*P*)], with 1,000 permutations and pathway size filters of 10–500 genes. The workflow also evaluated overlap between input genes and pathway databases to identify potential identifier mismatches and generated tabulated enrichment results together with summary plots of the top enriched pathways.

### In vitro cellular specificity assays

To assess oxidative stress associated with cytotoxicity in DPCs, cells were seeded in follicle dermal papilla cell growth medium with supplement (PromoCell) containing 1% penicillin-streptomycin at 1,000 cells in 20 µL per well in 384-well plates and allowed to attach for 12–14 h. Compounds were then added as indicated and incubated for 10 h, after which ROS levels were quantified using the ROS-Glo™ Assay (Promega) according to the manufacturer’s instructions. Skin sensitization signaling was additionally assessed using HepG2 cells engineered with an ARE-Nrf2 luciferase reporter. HepG2-ARE-Nrf2 cells were plated at 5,000 cells in 20 µL per well in 384-well plates, allowed to attach overnight, and treated with compounds for 24 h prior to measuring luciferase activity using the Bright-Glo™ Luciferase Assay System (Promega) following the manufacturer’s protocol. To evaluate potential genotoxicity, an SOS chromotest was performed in E. coli. Overnight *E. coli* cultures were grown in LB at 37 °C with shaking, diluted in fresh LB to an OD_600_ of 0.05–0.1, and dispensed into clear flat-bottom 384-well plates (20 µL per well). Compounds were added and incubated for 2 h at 37 °C. Cultures were then incubated with blue chromogen reagent (100 mM PBS, 0.1% Triton X-100, 1 mg/mL X-gal) for 1 h at 37 °C, and absorbance was measured at OD_600_ to assess SOS response activation. 4-nitroquinoline N-oxide (4-NQO) was used as a positive control.

### Macrophage activation assay

Primary human CD14^+^ monocytes were obtained from de-identified leukopaks (BioIVT) following PBMC enrichment in a Ficoll gradient. Isolated monocytes were subsequently differentiated into macrophages using the Immunocult-SF macrophage differentiation medium supplemented with M-CSF (Stem Cell Tech). Briefly, monocytes were plated in 96-well plates (NEST) at a concentration of 1×10^6^ cells/mL and differentiated according to the manufacturer’s protocol. The resulting unpolarized and matured macrophages were treated with lipopolysaccharide (LPS), NV-273, NV-623, NV-624, or NV-1065 for 24 h and their activation was assessed by flow-cytometric phenotyping. Gates were set for each sample based on fluorescence minus one (FMO) controls.

### Statistical analysis

Statistical analyses were performed using Prism v10.6.2 (GraphPad) and Python 3.12. Equality of variances was determined using Levene’s test. Normally distributed data were expressed as mean ± standard deviation.

## Supporting information

Representative flow cytometry gating

HPLC, MS, and NMR spectra of NV-623 and NV-624

## Reporting summary

Further information on research study design is available.

## Data availability

Data supporting the findings of the study are available within the main manuscript or Supplementary Information. All outstanding data requests may be made to the corresponding author.

## Acknowledgements

Z.Q., Y.L., S.E.C., L.D., Q.Y., L.T., G.Z., E.M.Z., and D.K.Y.Z. would like to thank the team at LK Innovations (https://lkinnovations.org/) and Pacagen, Inc. (https://pacagen.com/) for their continuous and unwavering support. Z.Q., Y.L., S.E.C, L.D, G.Z., E.M.Z, and D.K.Y.Z. would like to thank the investors of Pacagen, Inc. for their support. The authors thank the WuXi Research Service Chemistry group for their synthesis of NV-623 and NV-624 and NGS-Lab for performing the RNA sequencing experiments.

## Contributions

Z.Q., Y.L., S.E.C., F.W., E.M.Z., and D.K.Y.Z. conceived and designed the experiments. Z.Q., S.E.C., L.D., Q.Y., L.T., G.Z., A.L., S.O., and D.K.Y.Z. performed the experiments. Z.Q., Y.L., S.E.C., F.W., E.M.Z., and D.K.Y.Z. analyzed the data. Z.Q., Y.L., S.E.C., E.M.Z., and D.K.Y.Z. wrote the manuscript. All authors discussed the results and provided comments on the manuscript. The principal investigator is D.K.Y.Z.

## Competing interests

E.M.Z. and D.K.Y.Z. are co-founders of LK Innovations, and Z.Q., Y.L., S.E.C., L.D., Q.Y., L.T., G.Z. have equity interests in LK Innovations. F.W. is a co-founder of Integrated Biosciences, and A.L., S.O., and F.W. have equity interests in Integrated Biosciences.

## Supporting information

**Supplementary Table 1.** Donor information of primary cells used for functional studies.

|  | Age | Gender | Ethnicity |
| --- | --- | --- | --- |
| **DPC Donor 1** | 46 | F | Caucasian |
| **DPC Donor 2** | 45 | F | Caucasian |
| **DPC Donor 3** | 55 | F | Caucasian |
| **HEK Donor** | 28 | F | Caucasian |
| **HDMEC Donor** | 6 | M | Caucasian |
| **HDF Donor** | 66 | F | Caucasian |
| **PBMC Donor** | 60 | F | N/A |
| **HRCEpC Donor** | 76 | M | Caucasian |
| **HPAd Donor** | 21 | M | Caucasian |
| **HSkMC Donor** | 56 | M | Caucasian |

\*M: male; F: female; N/A: not available

**Supplementary Figure 1.**
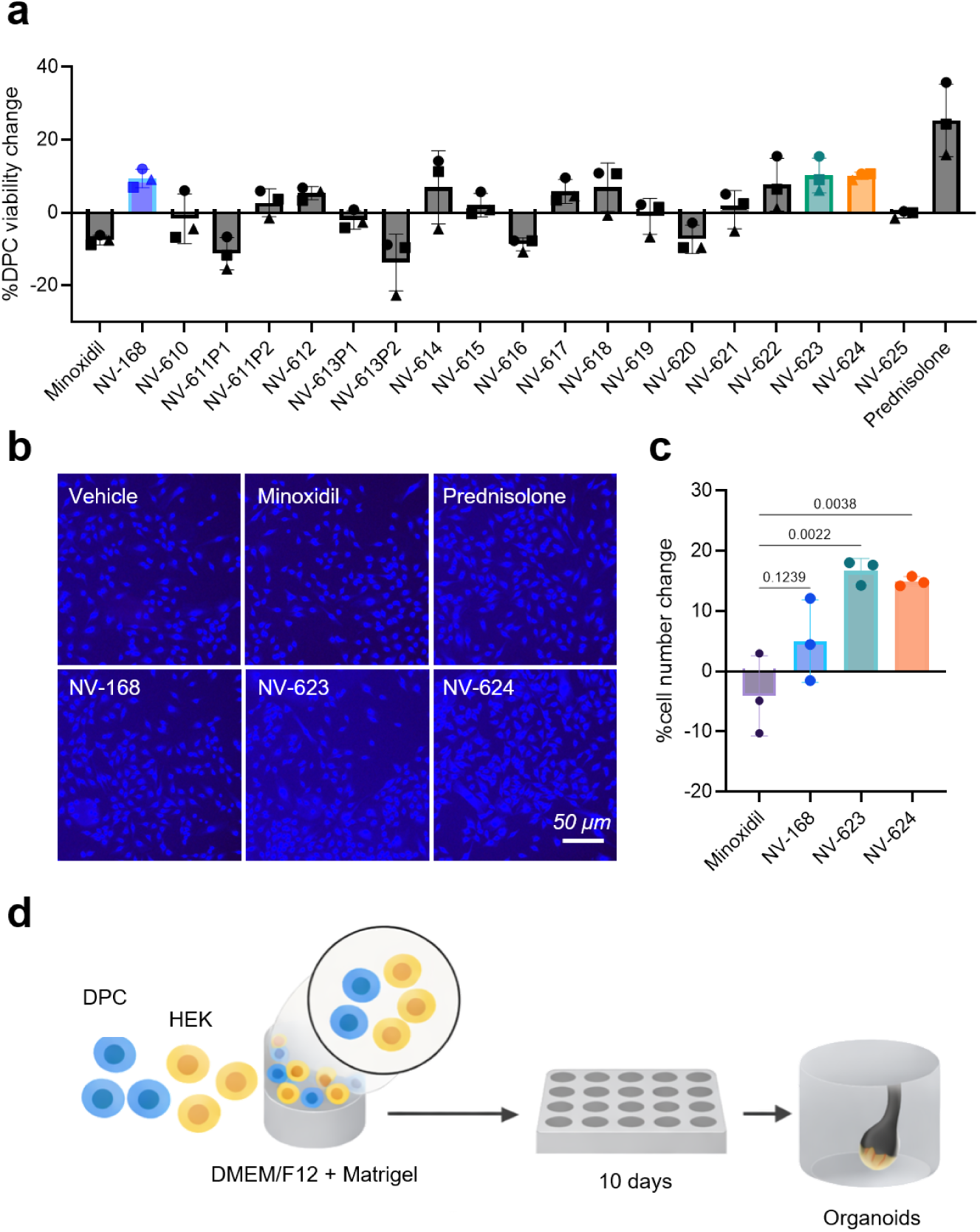
Cell counting of compound-treated DPCs and scheme of follicle organoid studies. (a) Overview of NV-168 derivative efficacy on DPC proliferation. (b) Representative images of cell cultures following treatment with lead compounds. The cells were fixed, stained with DAPI, and imaged. (c) Cell counts of DAPI-stained cells. An in-house analysis program was developed to analyze the images by detecting cell contours under high contrast conditions. (d) Hair follicle organoids were assembled by co-culturing 5,000 DPCs and 5,000 HEKs in advanced DMEM/F12 (supplemented with 1% P/S and 2% matrigel). Data represent mean ± s.d. of n=3 technical replicates from over 3 experimental replicates. Data in (a) are from three independent donors. Data in (b) and (c) are from donor 1.

**Supplementary Figure 2.**
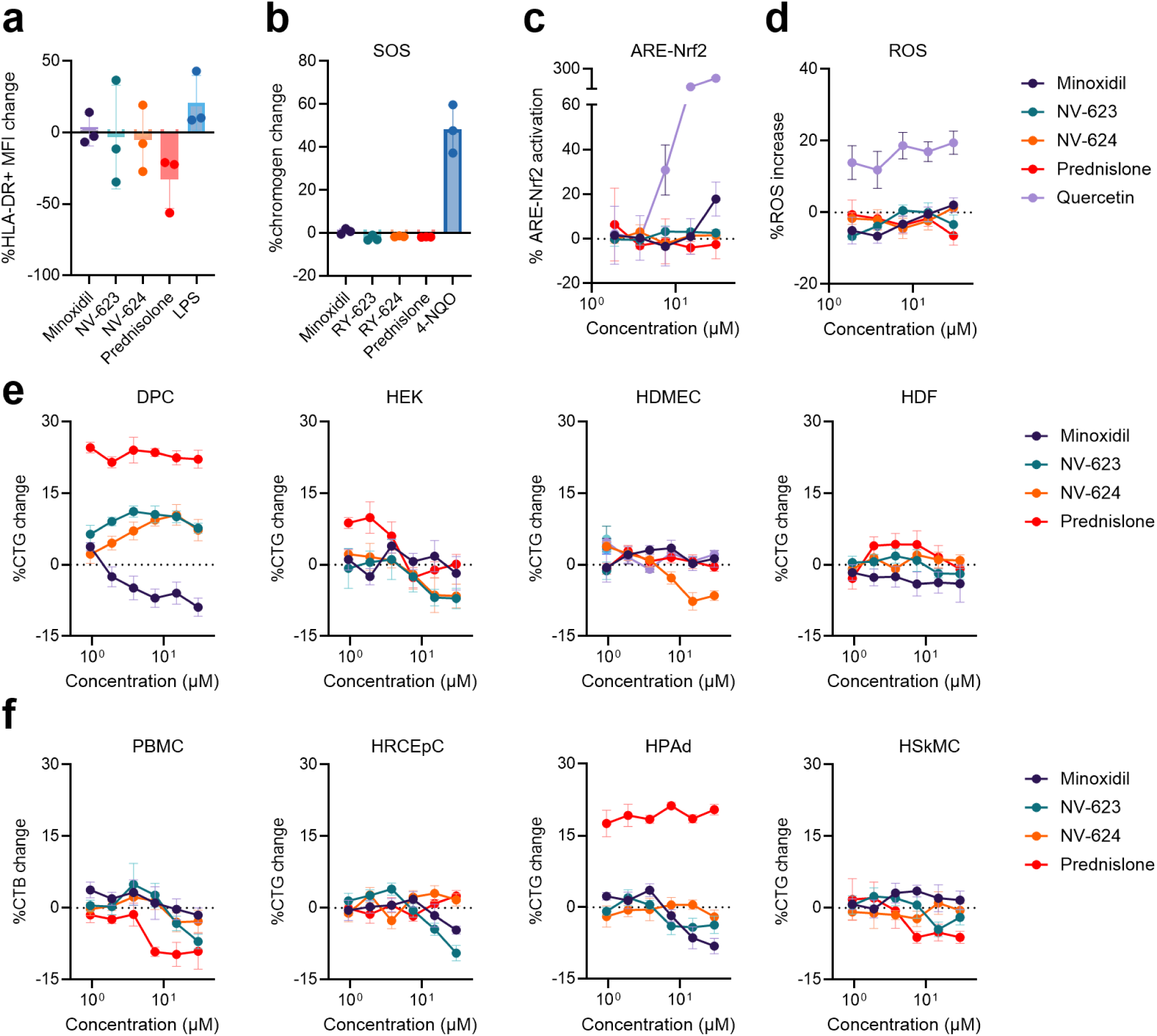
Cell specificity and safety evaluation of NV-623 and NV-624. (a) Activation of primary macrophages. (b) Genotoxicity measured by the SOS-chromotest. (c) Skin sensitization measured in engineered HepG2 ARE-Nrf2 reporter cells. (d) ROS production in primary DPCs. Effects on viability of (e) primary skin cells and (f) non-skin cells . Benchmark compounds are included for comparison. Data represent mean ± s.d. of n=3 technical replicates from over 3 experimental replicates.

**Supplementary Figure 3.**
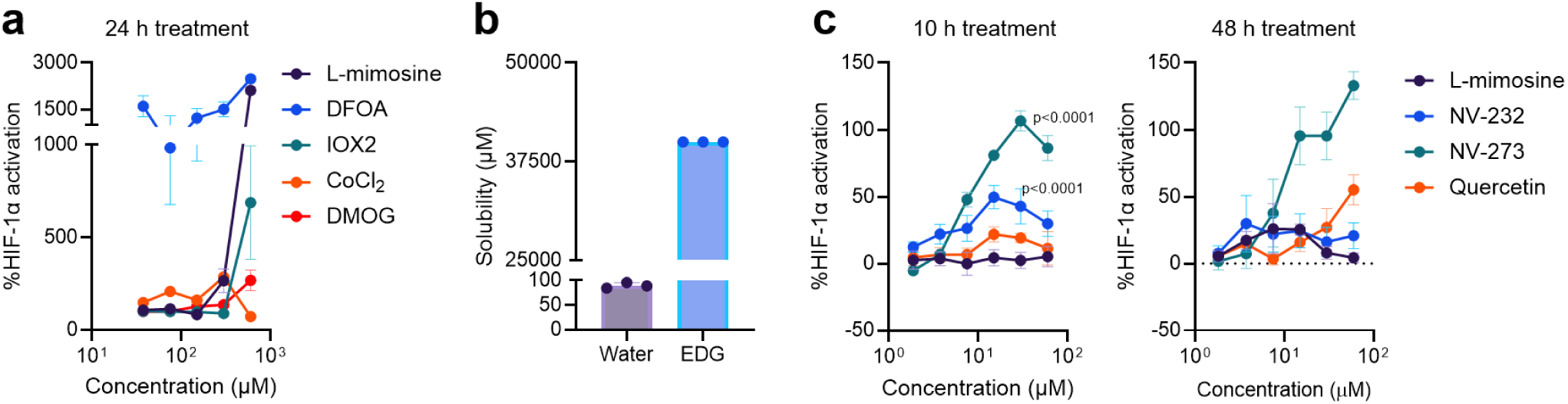
Time-dependent effects of PHD2 inhibitors. (a) HIF-1α activation of reported benchmark compounds including L-mimosine, DFOA, IOX2, cobalt chloride, and DMOG. (b) NV-273, despite moderate solubility in water, exhibited high solubility in ethoxydiglycol, supporting formulation-based solubility enhancement for improved solubility. DFOA: deferoxamine; DMOG: dimethyloxalylglycine; EDG: ethoxydiglycol. (c) HIF-1α stabilization after 10 h and 48 h treatment with NV-273, with the parent compound NV-232 and quercetin included for comparison. Data represent mean ± s.d. of n=3 technical replicates from over 3 experimental replicates.

**Supplementary Figure 4.**
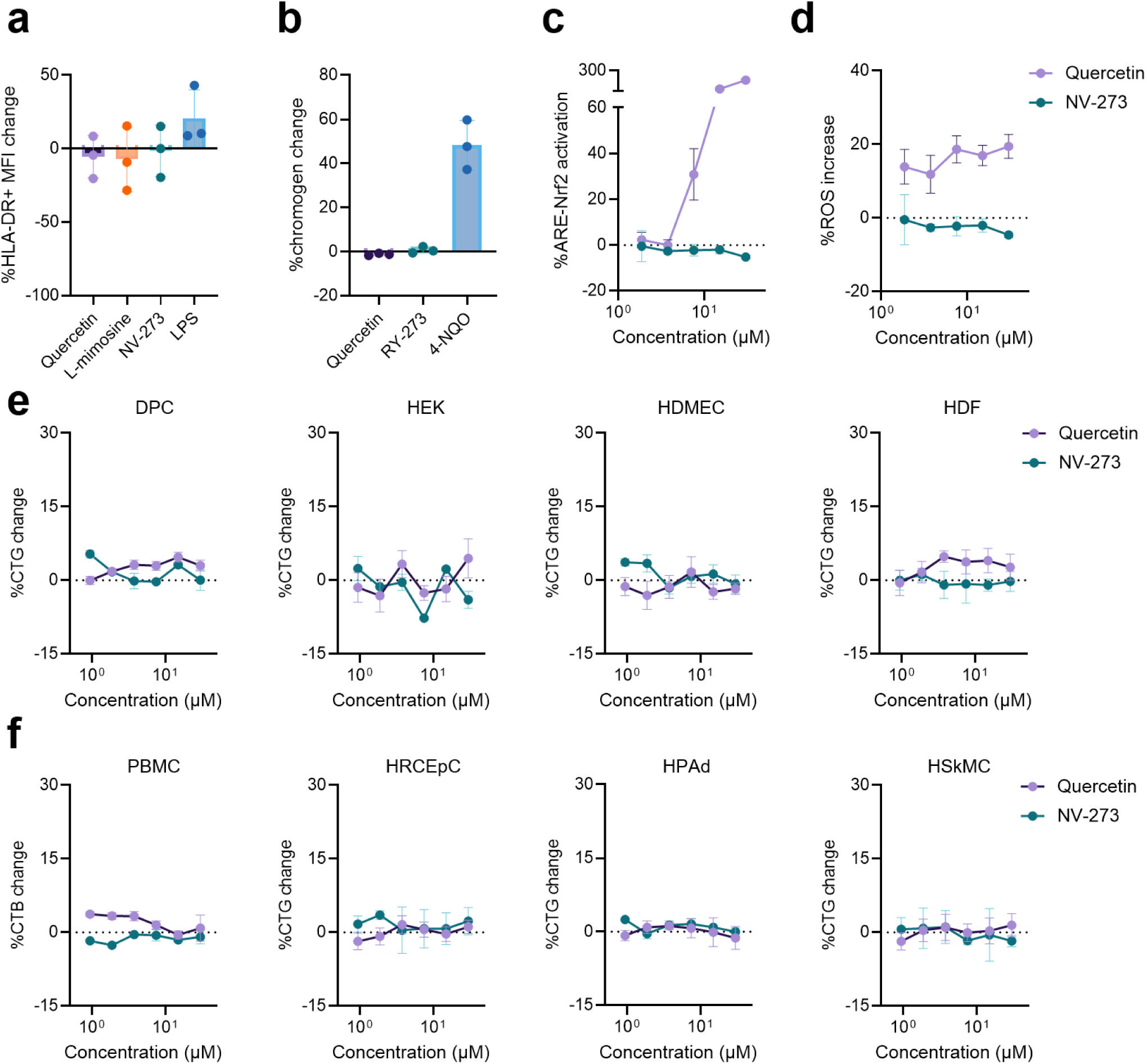
Cell specificity and safety evaluation of NV-273. (a) Activation of primary macrophages. (b) Genotoxicity measured by the SOS-chromotest. (c) Skin sensitization measured in engineered HepG2 ARE-Nrf2 reporter cells. (d) ROS production in primary DPCs. Effects on viability of (e) primary skin cells and (f) non-skin cells. Benchmark compounds are included for comparison. Data represent mean ± s.d. of n=3 technical replicates from over 3 experimental replicates.

**Supplementary Figure 5.**
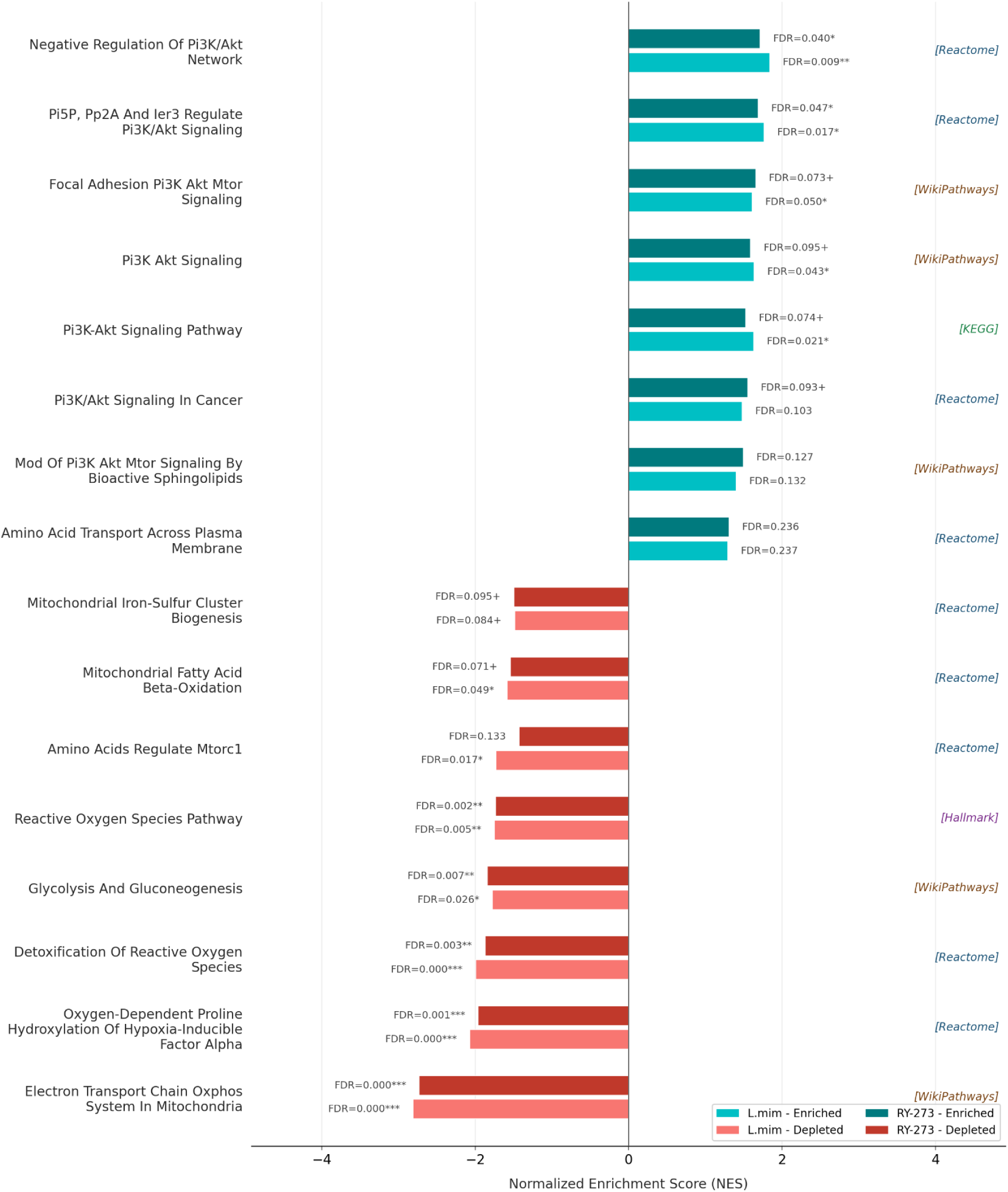
NV-273 elicited HIF-1α response comparable to L-mimosine. Positive NES values indicate enrichment and negative values indicate depletion. FDR values are indicated. Gene sets were derived from the Reactome, WikiPathways, KEGG and Hallmark collections.

**Supplementary Figure 6.**
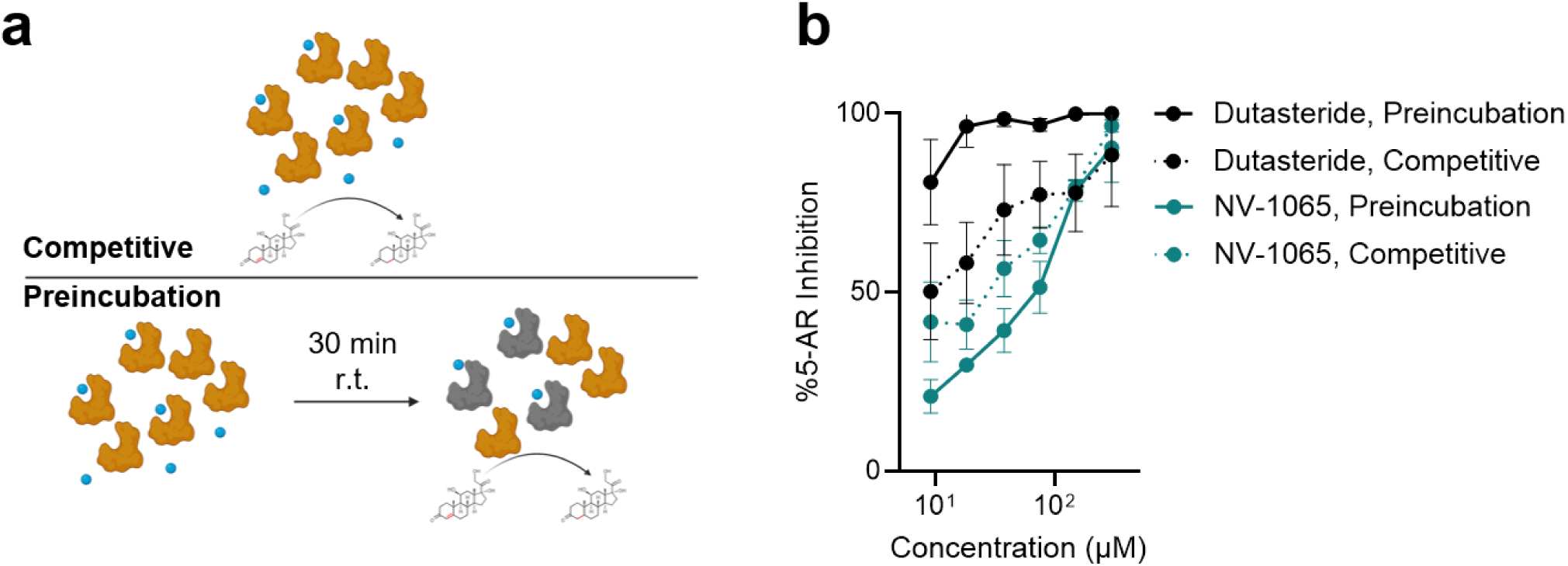
Time-dependent effects of 5-AR inhibitors. (a) Scheme of 5-AR inhibition measured under competitive conditions and with preincubation. (b) Effect of preincubation on 5-AR inhibition by NV-1065 and dutasteride. NV-1065 showed no significant change in potency, whereas dutasteride exhibited increased inhibition after preincubation, consistent with reversible inhibition by NV-1065. Data represent mean ± s.d. of n=3 technical replicates from over 3 experimental replicates.

**Supplementary Figure 7.**
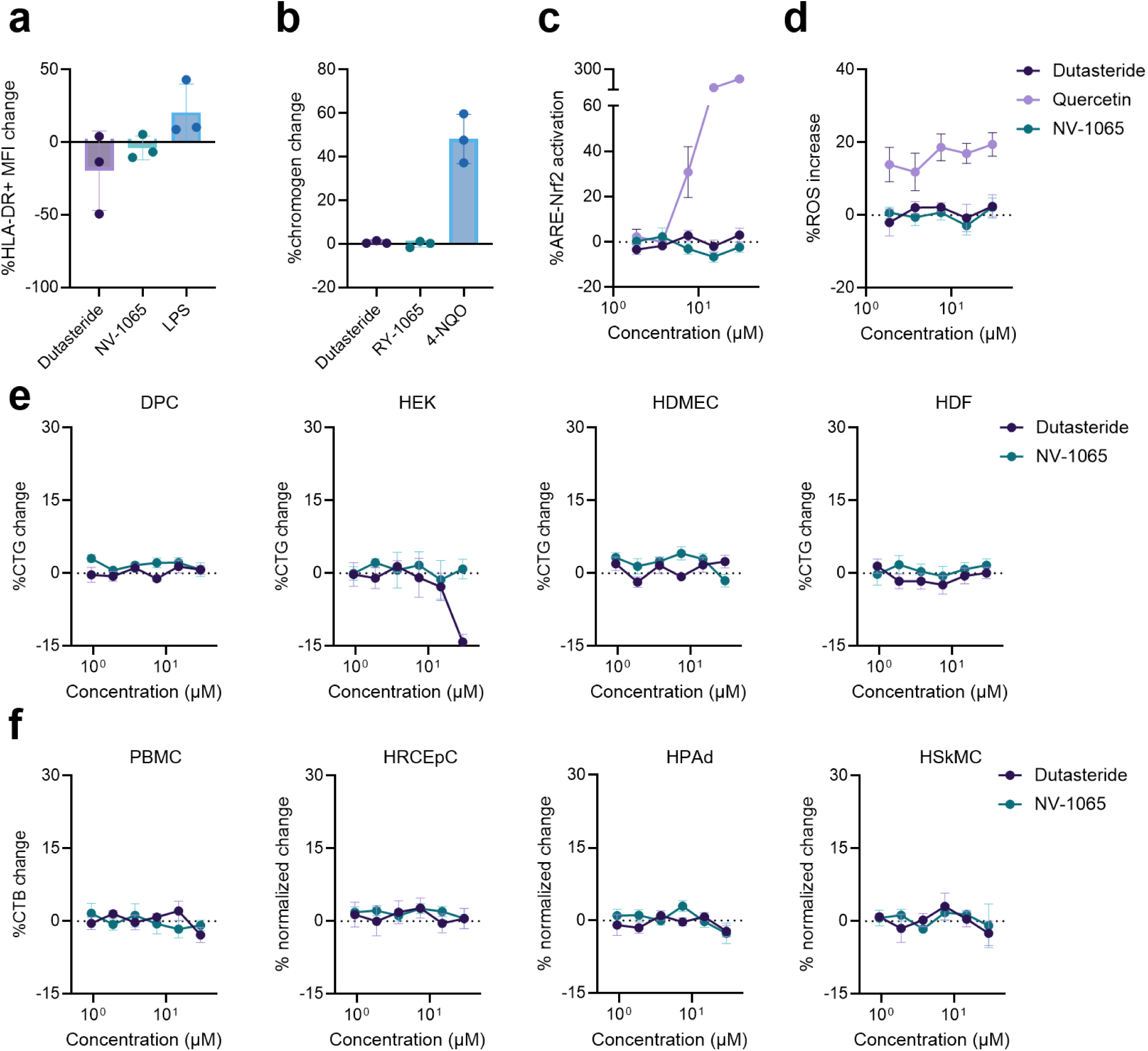
Cell specificity and safety evaluation of NV-1065. (a) Activation of primary macrophages. (b) Genotoxicity measured by the SOS-chromotest. (c) Skin sensitization measured in engineered HepG2 ARE-Nrf2 reporter cells. (d) ROS production in primary DPCs. (e,f) Effects on viability of (e) primary skin cells and (f) non-skin cells. Benchmark compounds are included for comparison. Data represent mean ± s.d. of n=3 technical replicates from over 3 experimental replicates.

**Supplementary Figure 8.**
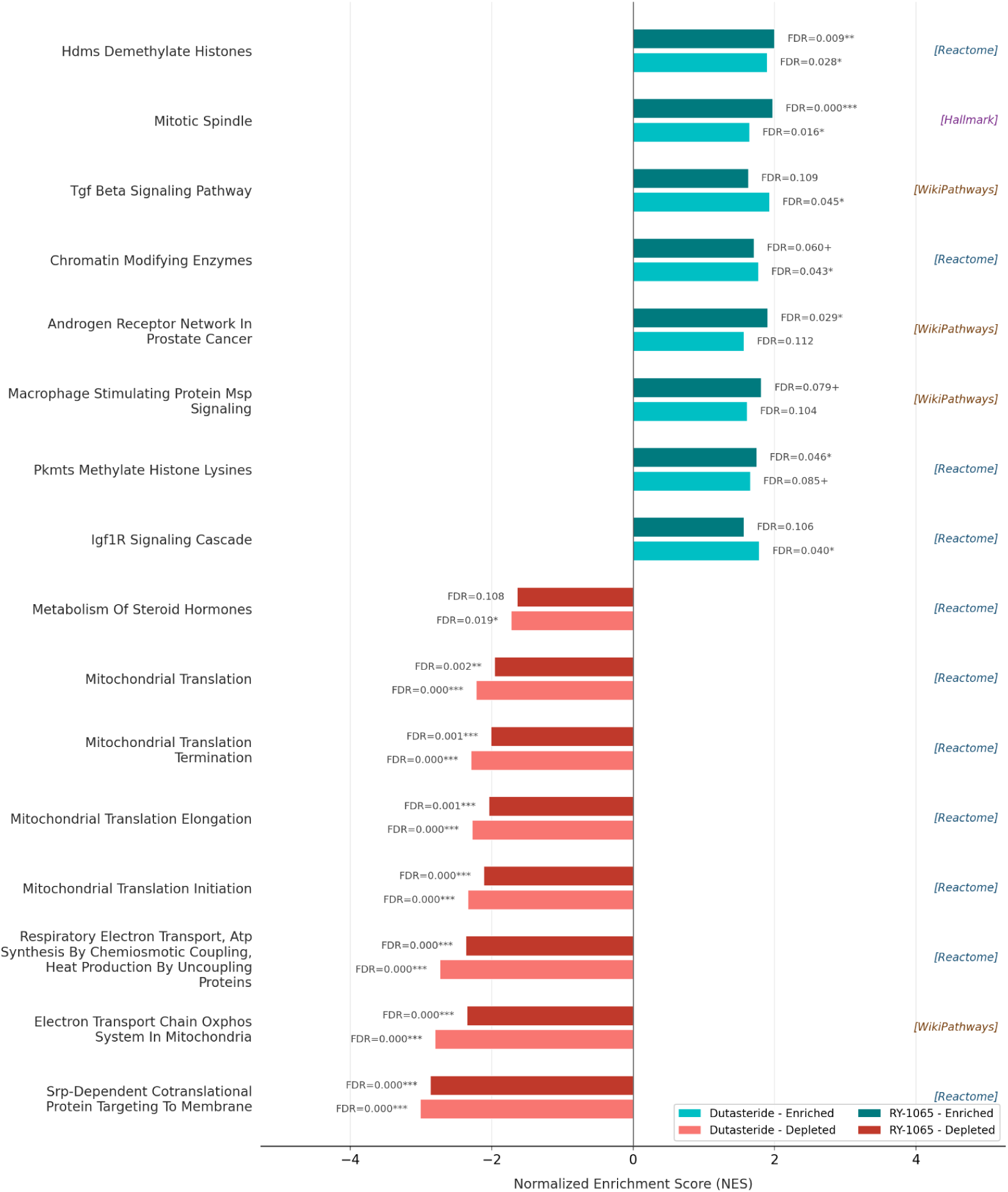
NV-1065 elicited AR-mediated responses comparable to dutasteride. Positive NES values indicate enrichment and negative values indicate depletion. FDR values are indicated. Gene sets were derived from the Reactome, WikiPathways, KEGG and Hallmark collections.

**Supplementary Figure 6.**
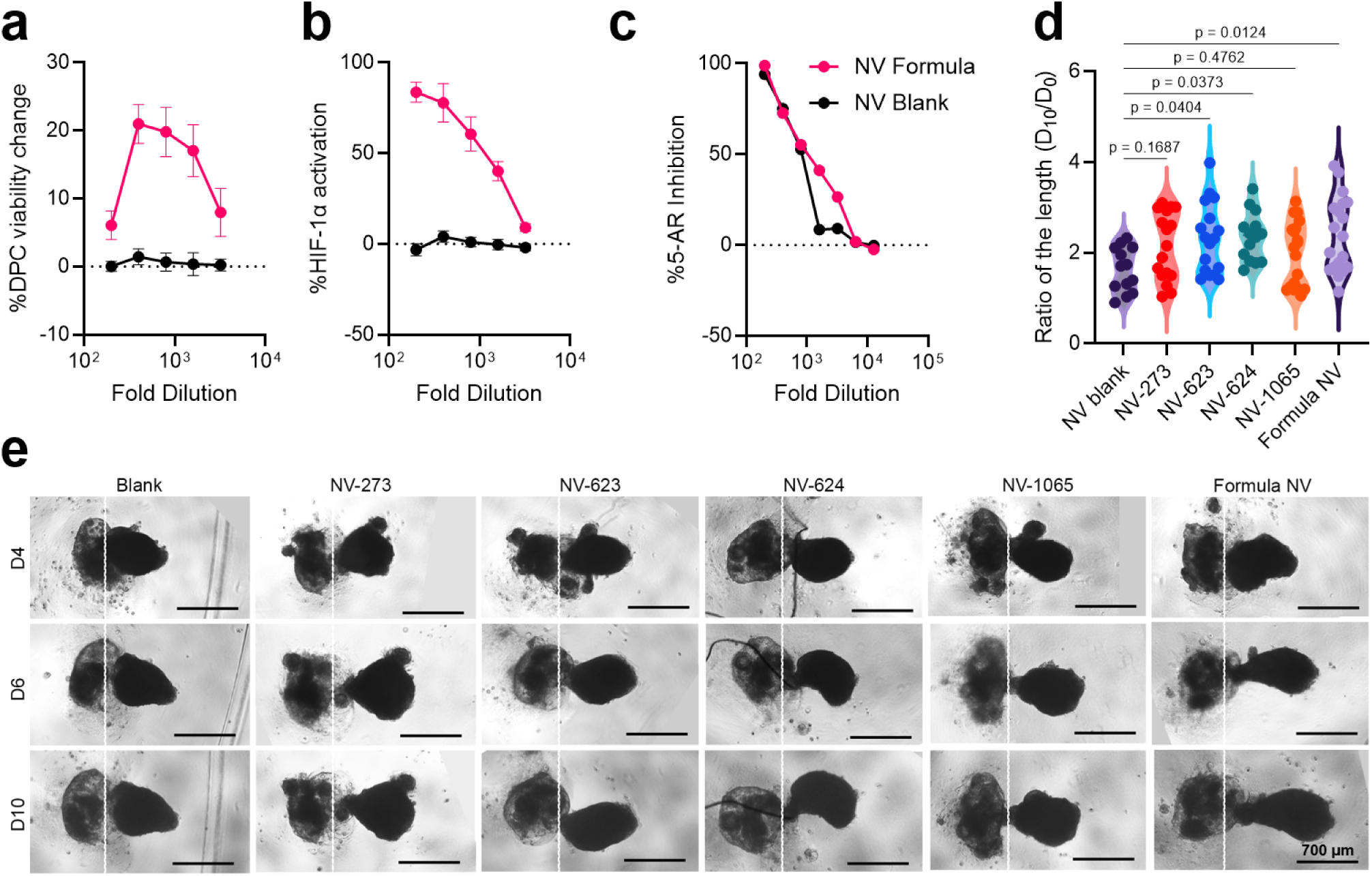
Characterization of cellular efficacy of formula NV. Dose-dependent responses of the formula NV and blank formulation in (a) DPC viability change, (b) HIF-1α activation, and (c) 5-AR inhibition. (d) quantitative analysis and (e) representative images showing combinatorial effects of formulated NV-273, NV-623, NV-624, and NV-1065 in promoting elongation of hair follicle organoids. All compounds were diluted to a final concentration of 60 μM. Data in (a-c) represent mean ± s.d. of n=3 technical replicates from 3 experimental replicates. Data in (d) represent n = 13 (blank), n = 17 (NV-273 and NV-623), n = 14 (NV-624), and n = 19 (NV-1065 and formula NV) from 3 experimental replicates. Images in (e) are representative of over 3 experimental replicates of 10 technical replicates.

