## Supplementary figures and images for "AI-enabled discovery of small molecules targeting complementary pathways for hair follicle rejuvenation"

### Representative flow cytometry gating

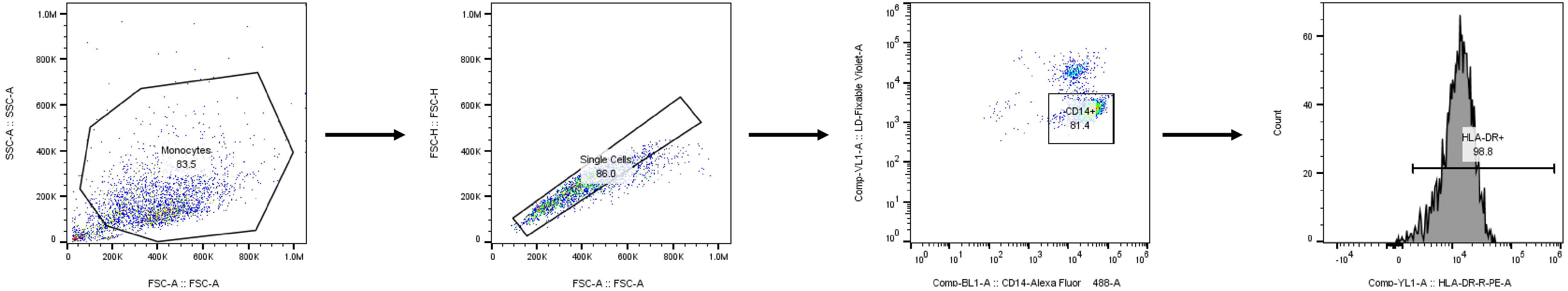

### Supplementary Fig. 1

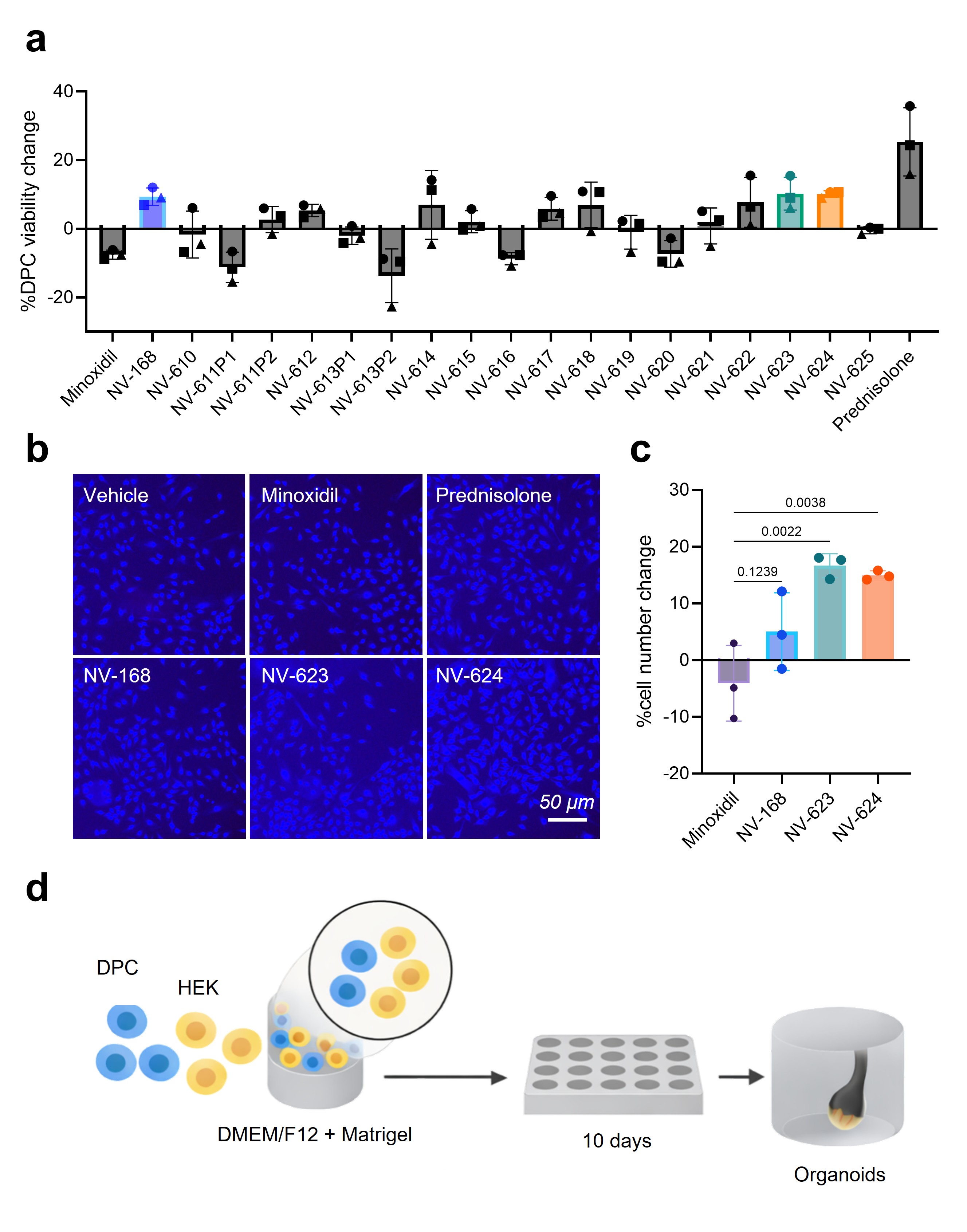

### Supplementary Fig. 2

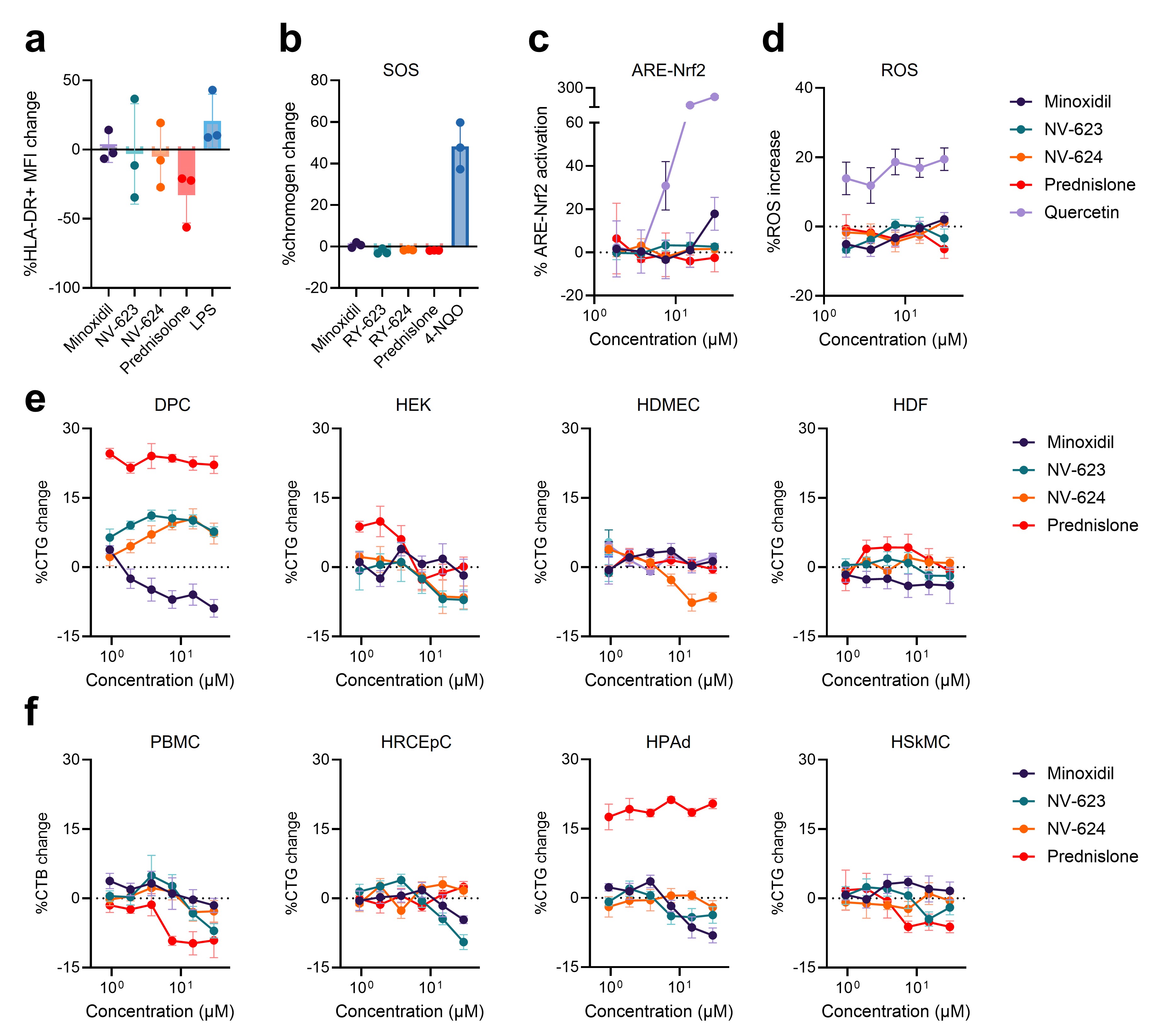

### Supplementary Fig. 3

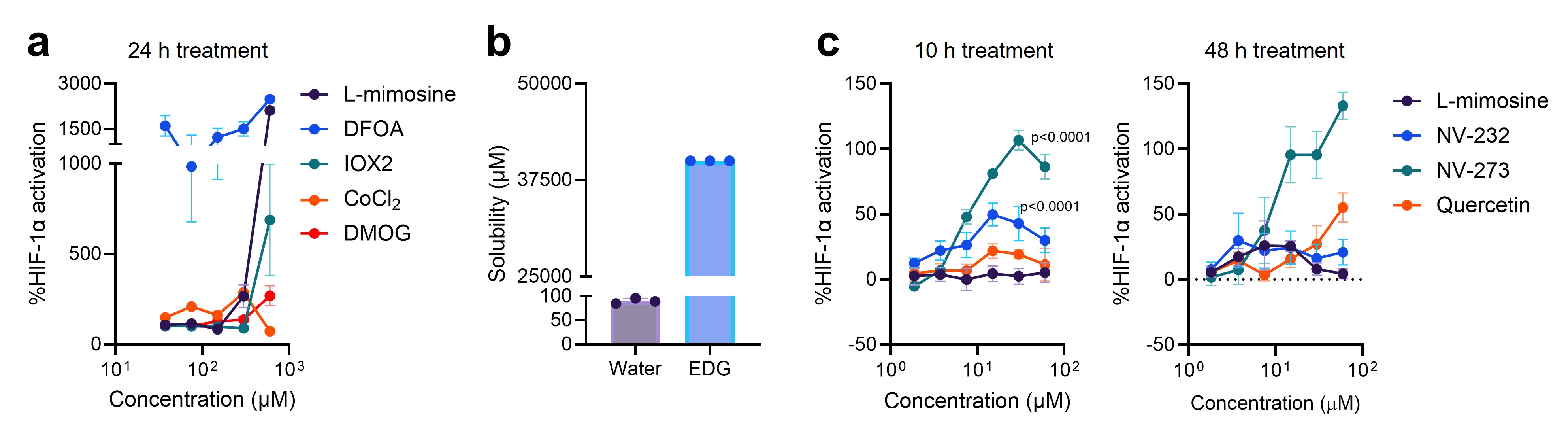

### Supplementary Fig. 4

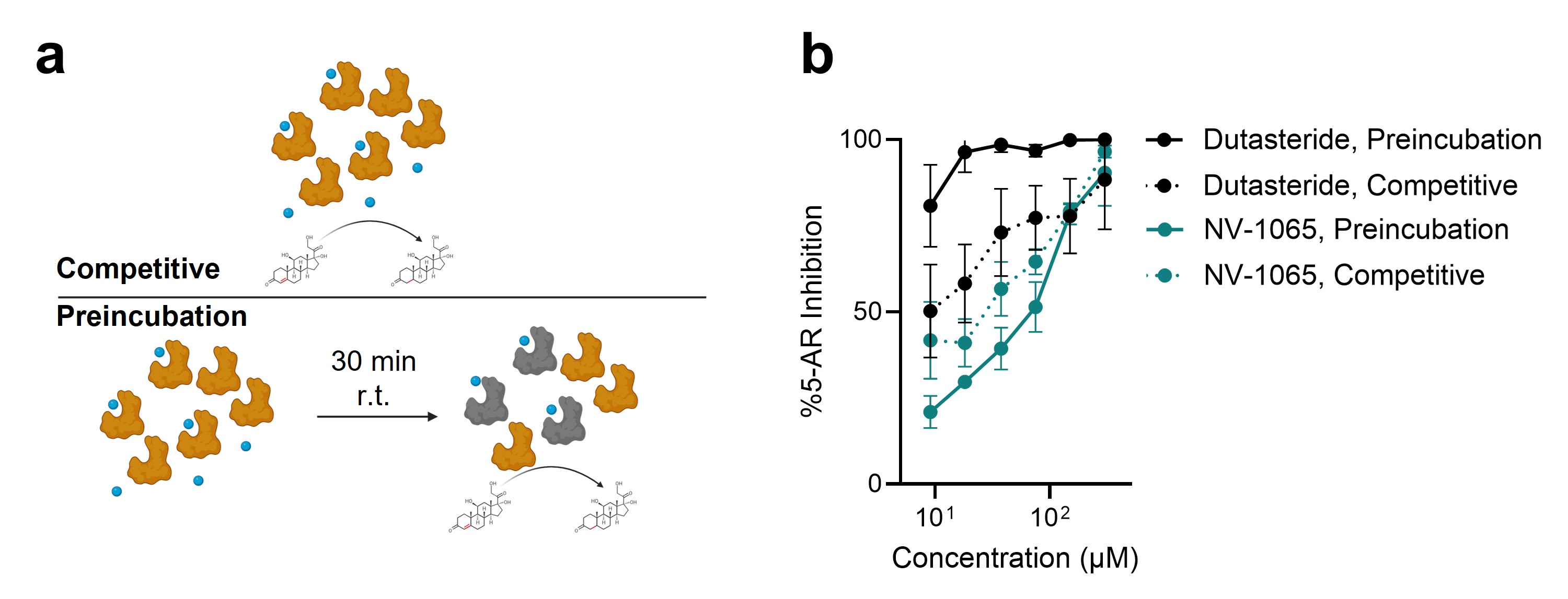

### Supplementary Fig. 5

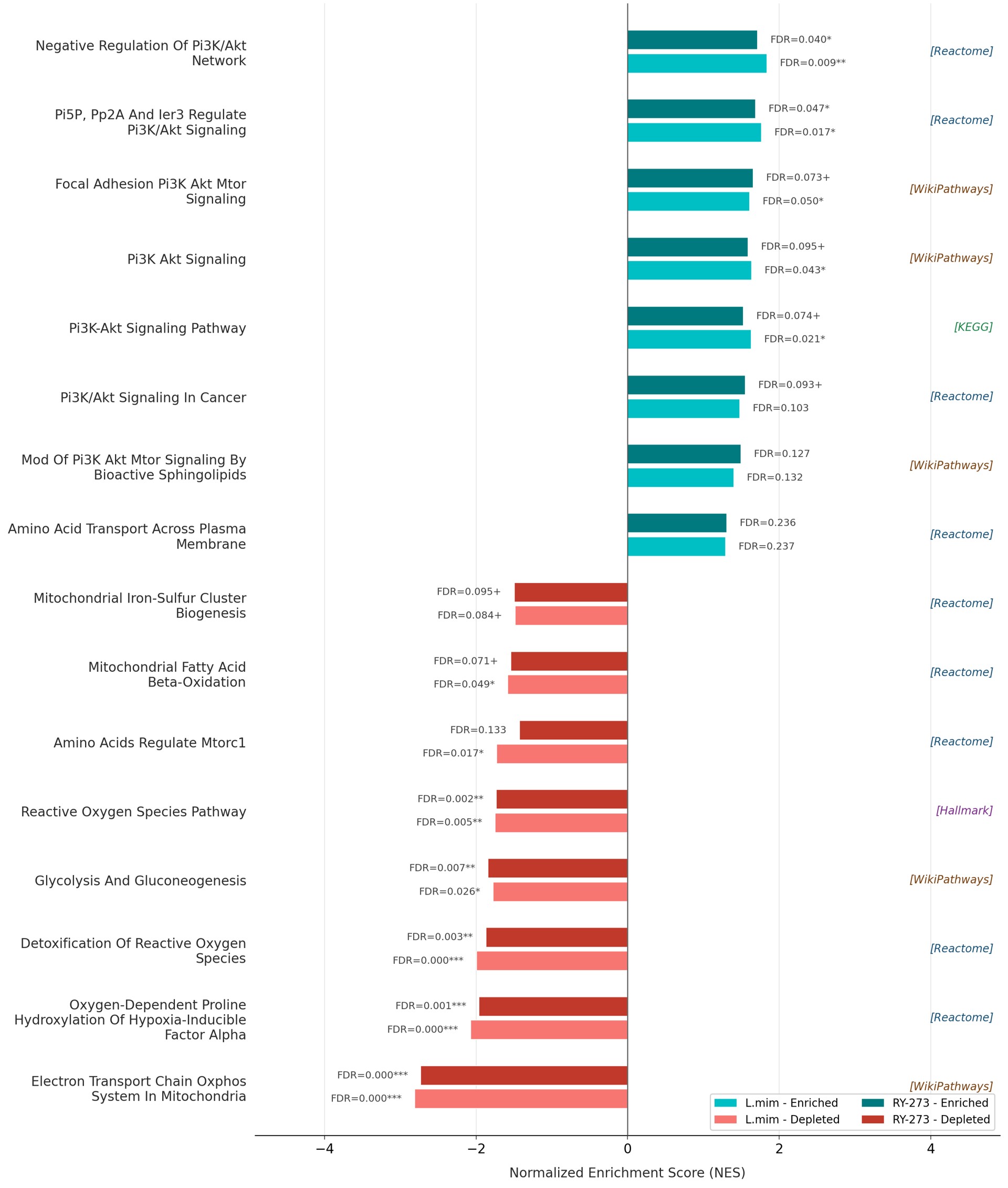

### Supplementary Fig. 6

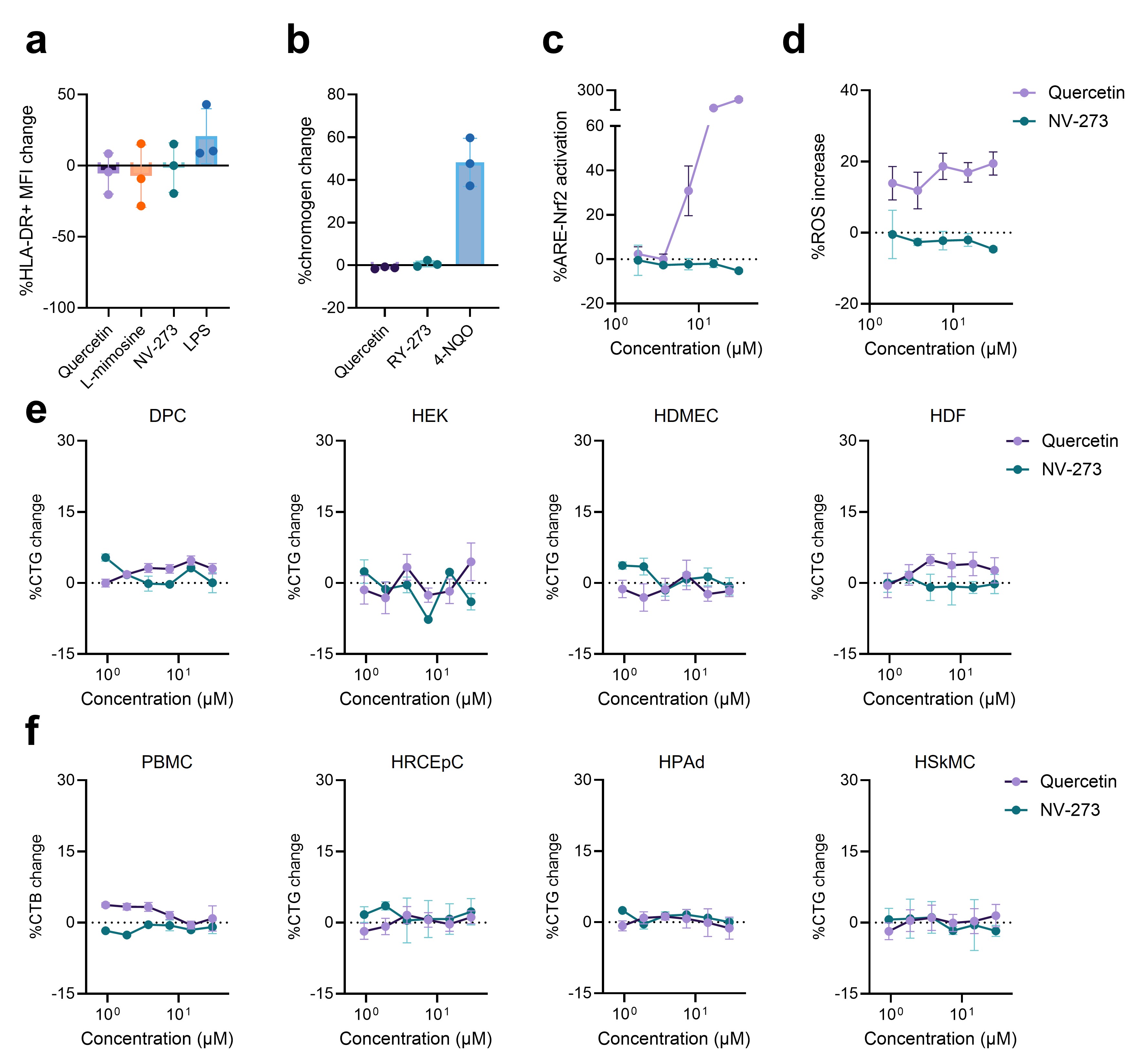

### Supplementary Fig. 7

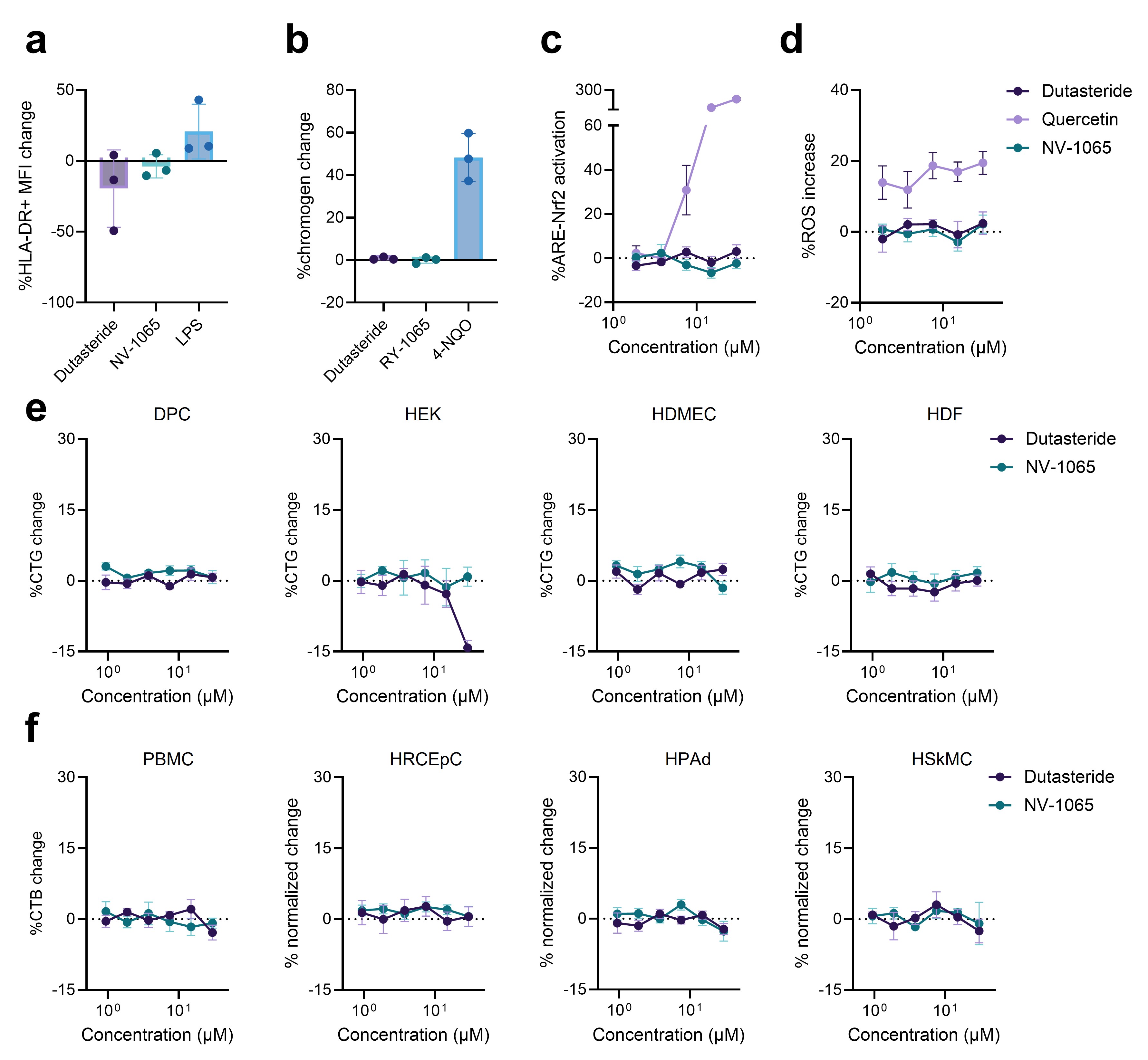

### Supplementary Fig. 8

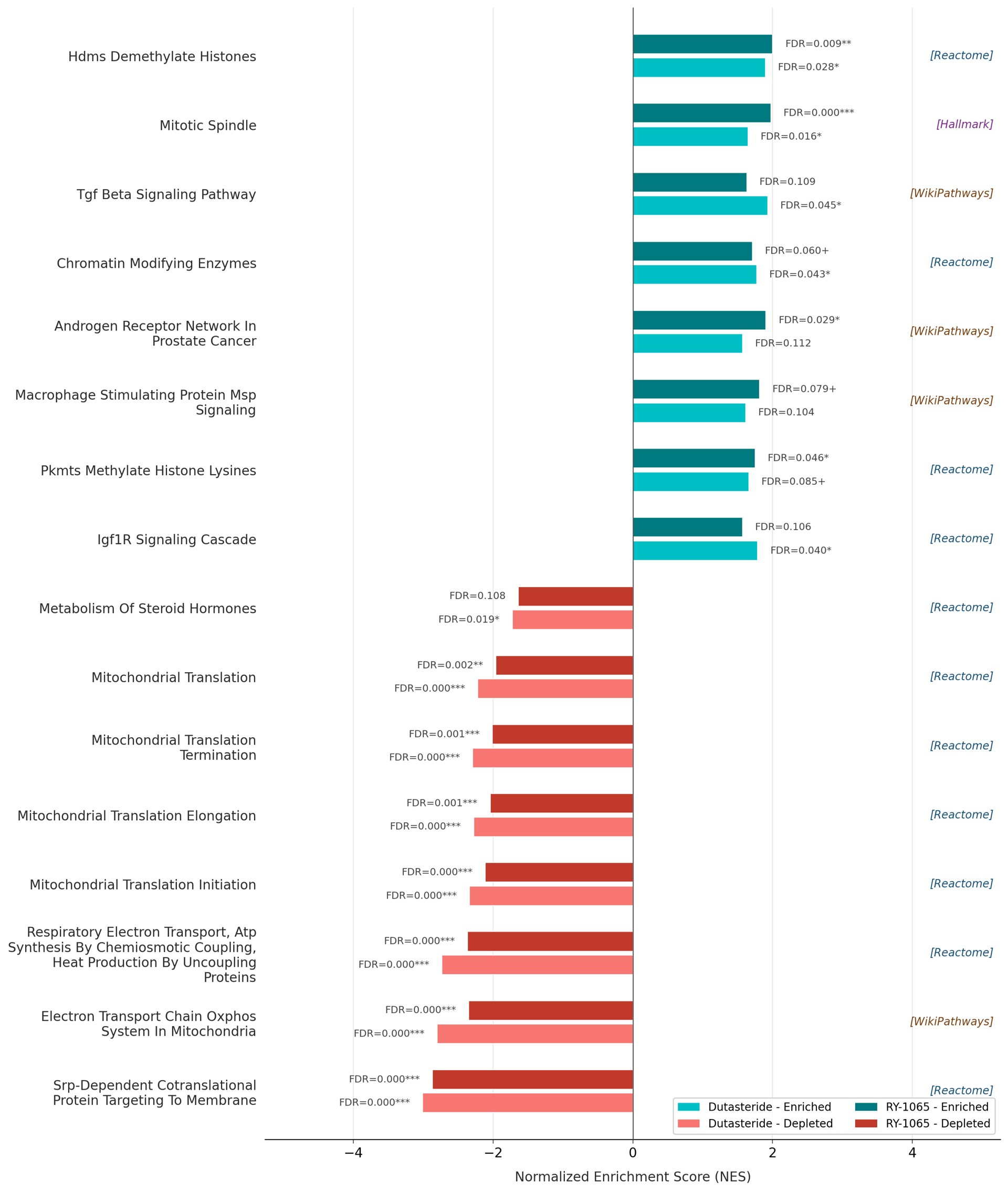

### Supplementary Fig. 9

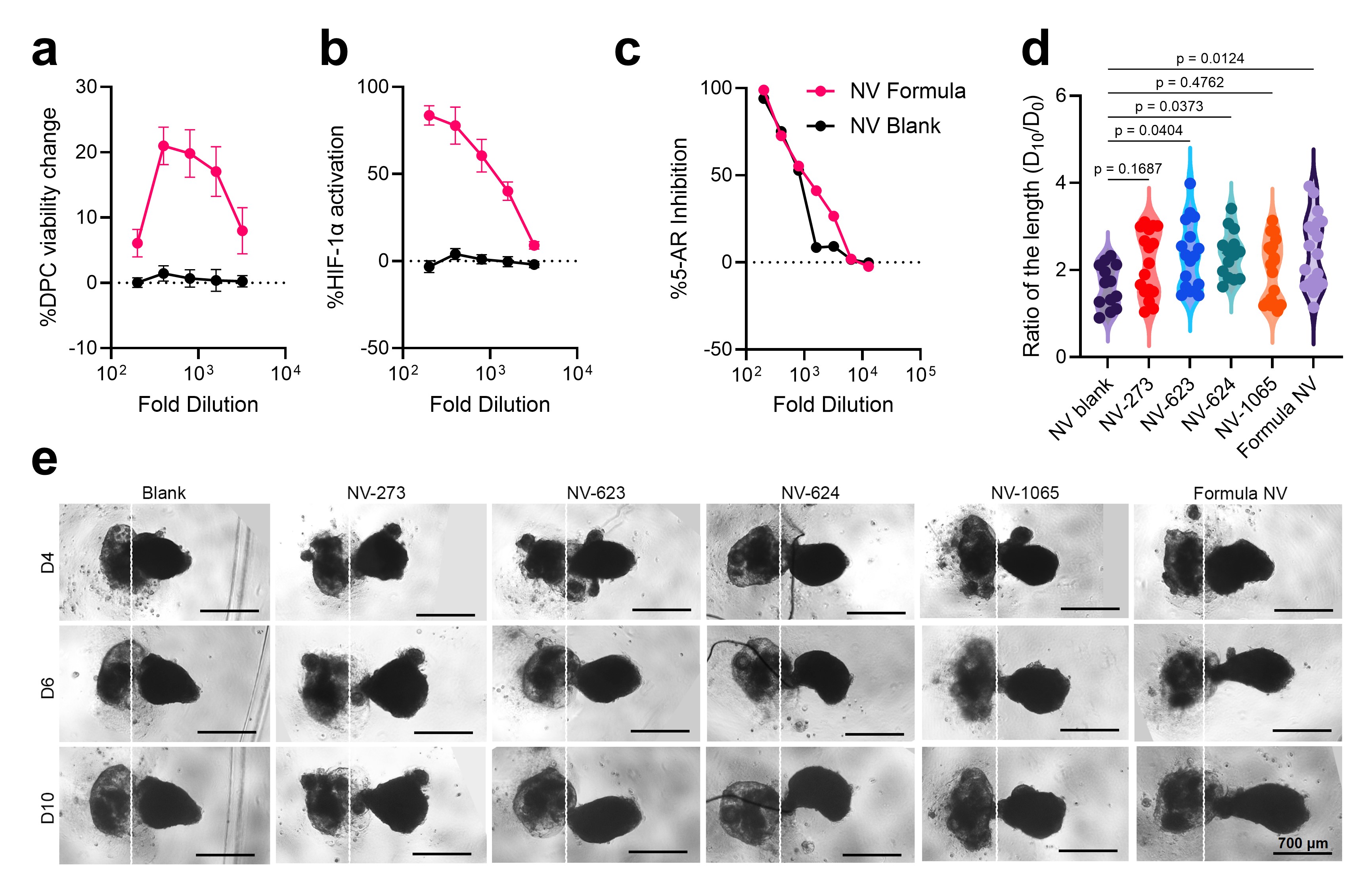
