## Supplementary material for "AI-enabled discovery of small molecules targeting complementary pathways for hair follicle rejuvenation": HPLC, MS, and NMR spectra of NV-623 and NV-624

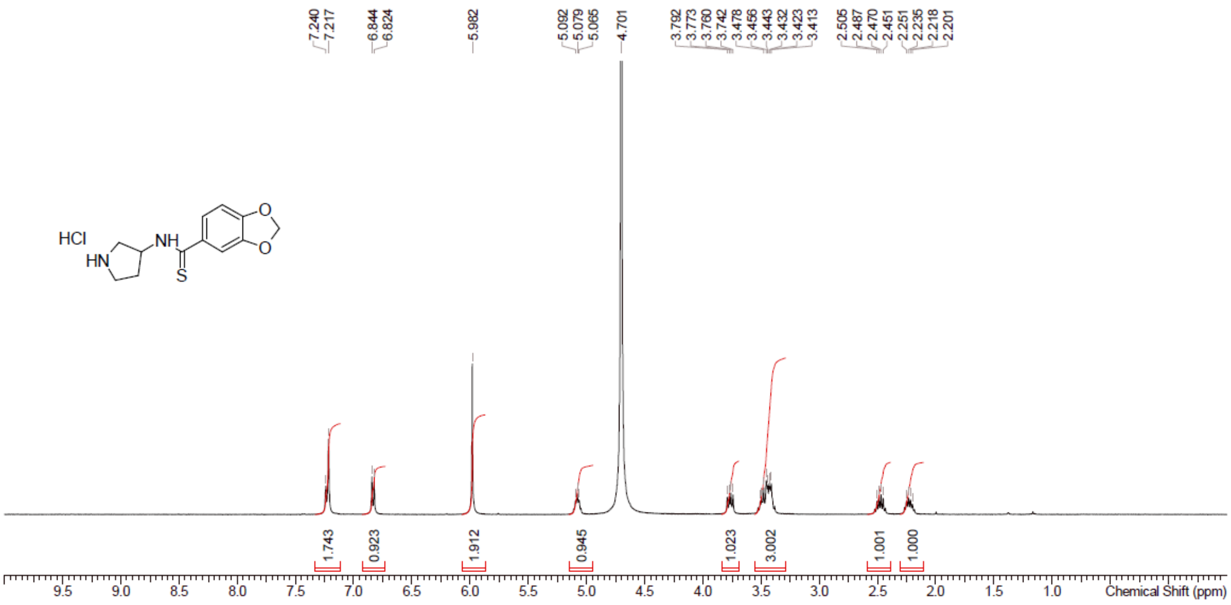


Supplementary Figure 10. [1]H NMR spectrum of RY-623.

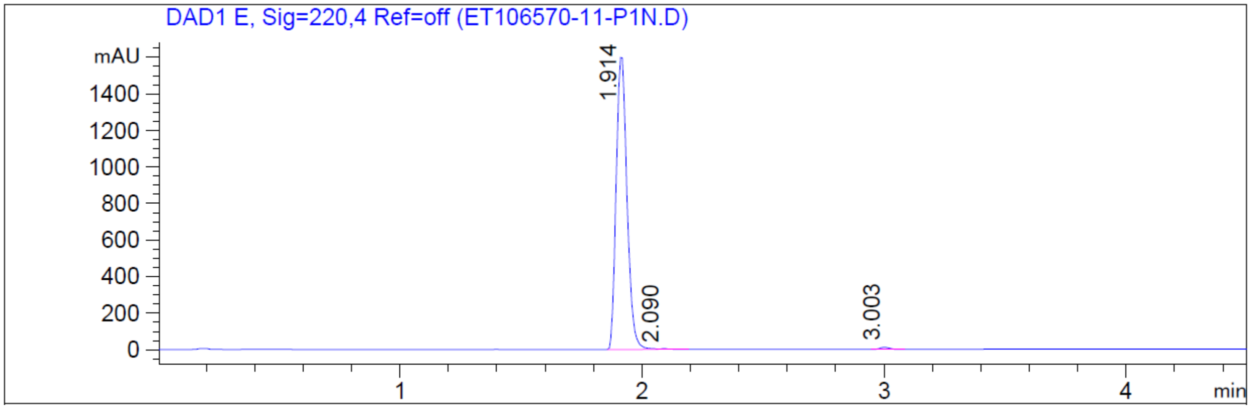


Supplementary Figure 11. HPLC spectrum of RY-623.

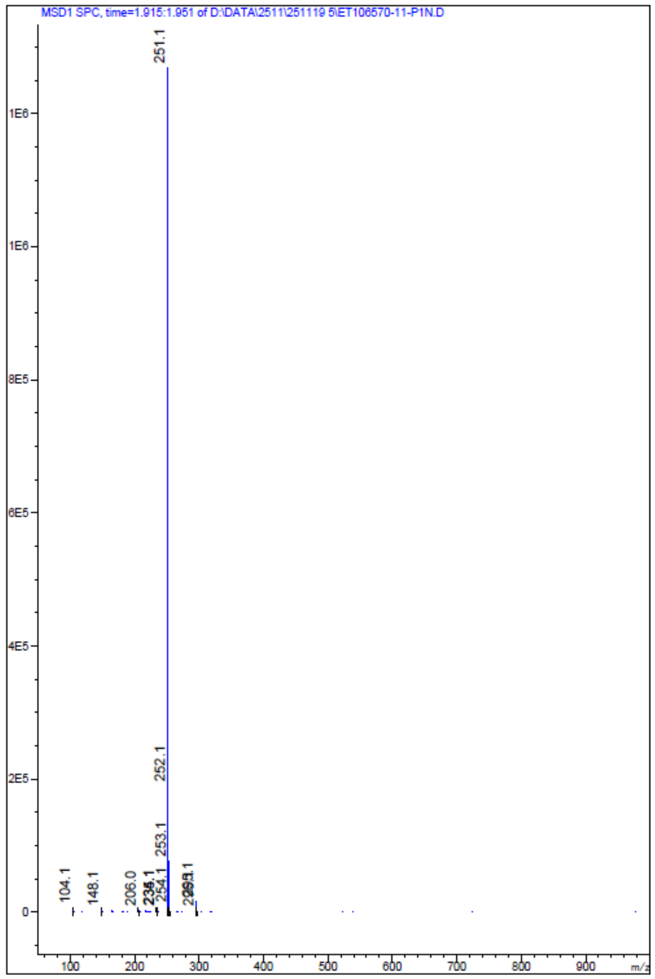


Supplementary Figure 12. MS spectrum of RY-623.

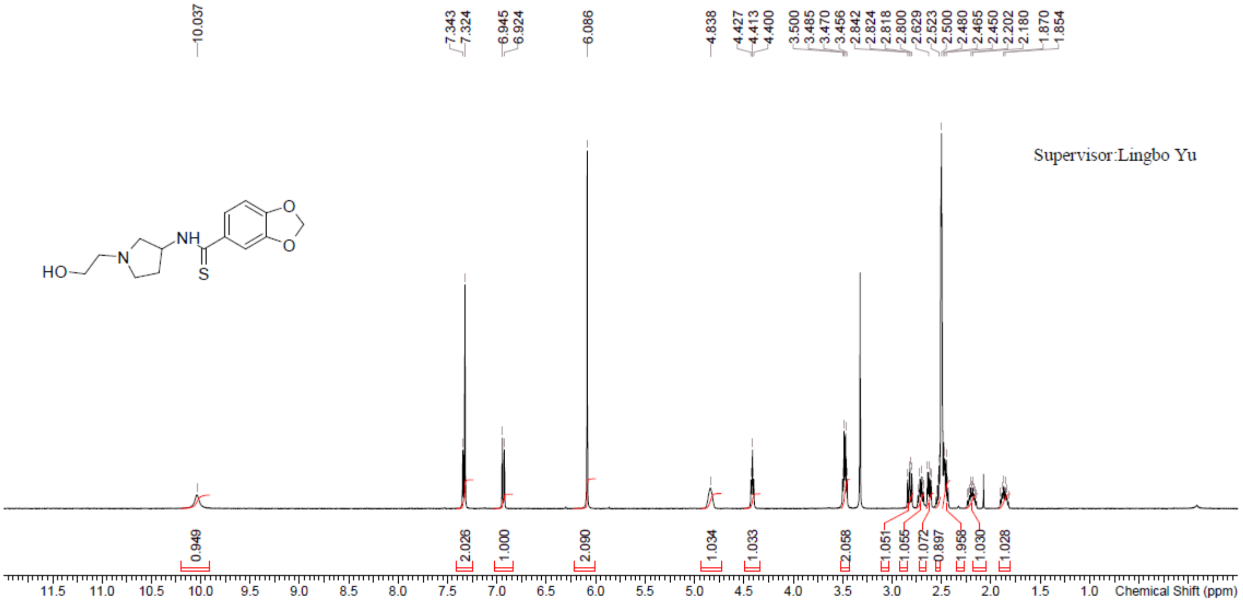

Supplementary Figure 13. $^{1}H$ NMR spectrum of RY-624.

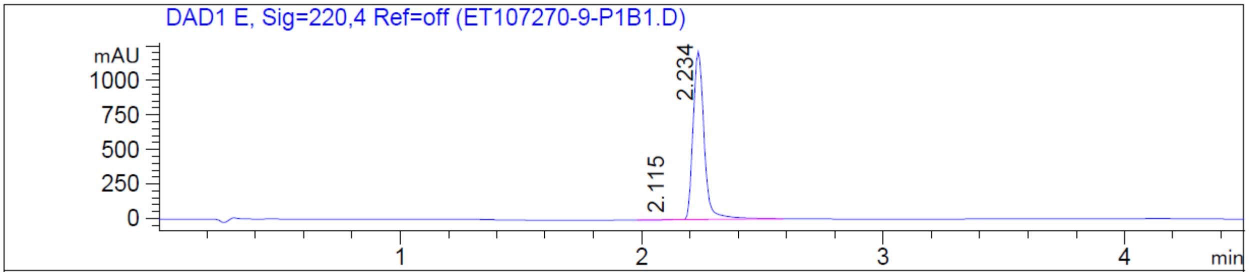

Supplementary Figure 14. HPLC spectrum of RY-624.

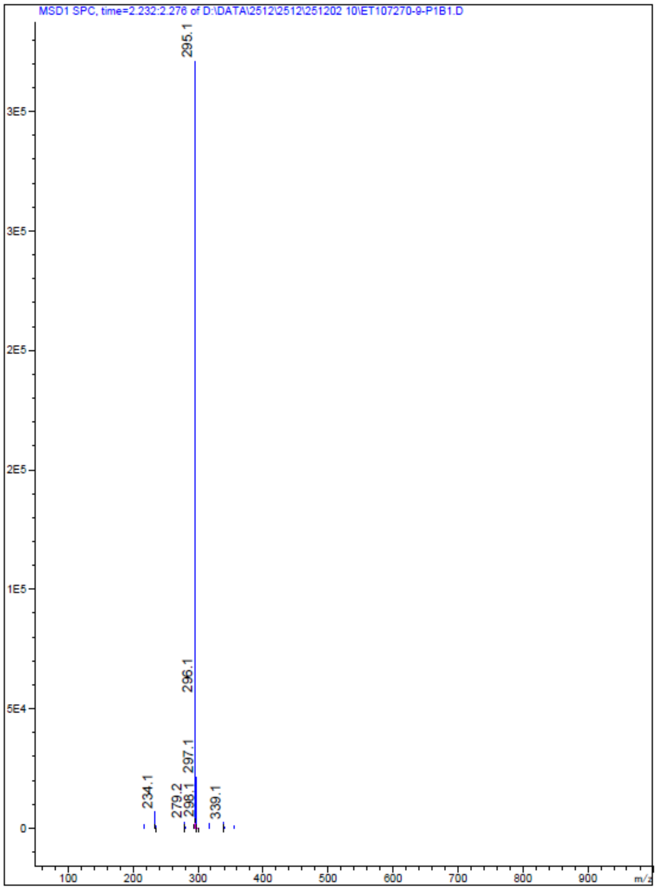

Supplementary Figure 15. MS spectrum of RY-624.
